# Mapping pediatric brain tumors to their origins in the developing cerebellum

**DOI:** 10.1101/2021.12.19.473154

**Authors:** Konstantin Okonechnikov, Piyush Joshi, Mari Sepp, Kevin Leiss, Ioannis Sarropoulos, Florent Murat, Martin Sill, Pengbo Beck, Kenneth Chun-Ho Chan, Andrey Korshunov, Felix Sahm, Maximilian Y. Deng, Dominik Sturm, John DeSisto, Andrew M. Donson, Nicholas K. Foreman, Adam L. Green, Giles Robinson, Brent A. Orr, Qingsong Gao, Emily Darrow, Jennifer L. Hadley, Paul A. Northcott, Johannes Gojo, Marina Ryzhova, Daisuke Kawauchi, Volker Hovestadt, Mariella G. Filbin, Andreas von Deimling, Marc Zuckermann, Kristian W. Pajtler, Marcel Kool, David T.W. Jones, Natalie Jäger, Lena M. Kutscher, Henrik Kaessmann, Stefan M. Pfister

**Affiliations:** Hopp Children’s Cancer Center (KiTZ), Heidelberg, Germany; Division of Pediatric Neurooncology, German Cancer Research Center (DKFZ) and German Cancer Consortium (DKTK), Heidelberg, Germany; Center for Molecular Biology of Heidelberg University (ZMBH), DKFZ-ZMBH Alliance, Heidelberg, Germany; Clinical Cooperation Unit Neuropathology, German Cancer Research Center (DKFZ), Heidelberg, Germany; Department of Neuropathology, Institute of Pathology, Heidelberg University Hospital, Heidelberg, Germany; Division of Pediatric Glioma Research, German Cancer Research Center (DKFZ), Heidelberg, Germany; Department of Pediatric Hematology and Oncology, Heidelberg University Hospital, Heidelberg, Germany; Morgan Adams Foundation Pediatric Brain Tumor Research Program, University of Colorado School of Medicine, Aurora, CO, USA; Children’s Hospital Colorado, Aurora, CO, USA; Department of Oncology, St Jude Children’s Research Hospital, Memphis, TN, USA; Department of Pathology, St Jude Children’s Research Hospital, Memphis, TN, USA; Department of Developmental Neurobiology, St Jude Children’s Research Hospital, Memphis, TN, USA; Department of Pediatrics and Adolescent Medicine, Comprehensive Center for Pediatrics and Comprehensive Cancer Center, Medical University of Vienna, 1090 Vienna, Austria; Department of Neuropathology, NN Burdenko Neurosurgical Institute, Moscow, Russia; Department of Biochemistry and Cellular Biology, National Institute of Neuroscience, NCNP, Tokyo, Japan; Department of Pediatric Oncology, Dana-Farber Boston Children’s Cancer and Blood Disorders Center, Boston, USA; Broad Institute of Harvard and MIT, Cambridge, USA; Princess Máxima Center for Pediatric Oncology, 3584 CS Utrecht, the Netherlands; Developmental Origins of Pediatric Cancer Group, German Cancer Research Center (DKFZ), Heidelberg, 19. Germany

## Abstract

Understanding the cellular origins of childhood brain tumors is key for discovering novel tumor-specific therapeutic targets. Previous strategies mapping cellular origins typically involved comparing human tumors to murine embryonal tissues^1,2^, a potentially imperfect approach due to spatio-temporal gene expression differences between species^3^. Here we use an unprecedented single-nucleus atlas of the developing human cerebellum (Sepp, Leiss, et al) and extensive bulk and single-cell transcriptome tumor data to map their cellular origins with focus on three most common pediatric brain tumors – pilocytic astrocytoma, ependymoma, and medulloblastoma. Using custom bioinformatics approaches, we postulate the astroglial and glial lineages as the origins for posterior fossa ependymomas and radiation-induced gliomas (secondary tumors after medulloblastoma treatment), respectively. Moreover, we confirm that SHH, Group3 and Group4 medulloblastomas stem from granule cell/unipolar brush cell lineages, whereas we propose pilocytic astrocytoma to originate from the oligodendrocyte lineage. We also identify genes shared between the cerebellar lineage of origin and corresponding tumors, and genes that are tumor specific; both gene sets represent promising therapeutic targets. As a common feature among most cerebellar tumors, we observed compositional heterogeneity in terms of similarity to normal cells, suggesting that tumors arise from or differentiate into multiple points along the cerebellar “lineage of origin”.

---

Pediatric central nervous system (CNS) tumors represent one of the most fatal disease entities in children^4^. Despite major advances in the classification and diagnosis of these tumors, such as DNA methylation analysis^5^ and routine next-generation sequencing-based work-up^6^, contemporary treatment stratification does not match the tremendous inter-tumor heterogeneity observed in pediatric CNS tumors. Current radiation and chemotherapy approaches frequently result in neurocognitive disorders and life-long side effects, including secondary malignancies in patients surviving their primary disease^7^. Future therapeutic approaches must evolve to target tumor-specific vulnerabilities^8^, ideally without affecting normal tissue architecture – a difficult task, owing to the compositional and functional complexity of the brain^9^. To address these challenges, we focused on the cerebellum, the most frequent anatomic location of pediatric brain tumors^1,10^. We sought to uncover human cerebellar cellular diversity, identify cells vulnerable to tumor formation, and detect regulatory genes or surface markers as potential tumor specific therapeutic targets.

In this study, we concentrated on the three most common cerebellar tumor types in children: medulloblastoma (MB), posterior fossa ependymoma (PFA-EPD), and pilocytic astrocytoma (PA). Medulloblastoma, the most prevalent malignant cerebellar tumor, is classified into WNT, SHH, Group 3 and Group 4 molecular groups, and associated subgroups^11^. While WNT MBs are thought to arise from the lower rhombic lop in the dorsal brainstem rather than the cerebellum^12^, SHH MBs are known to originate from granule cell precursors^13,14^, whereas Group 4 MBs most likely arise from unipolar brush cells^1,2^. The cellular origin of Group 3 MBs is still less clearly pinpointed. PFA, a hindbrain-specific ependymoma type^15^, is frequently fatal due to its chemotherapy resistance. In previous studies, this tumor class showed closest similarities to radial glial cell subtypes^16^ – “roof-plate-like stem cells” and “gliogenic progenitors” – when comparing it to murine cerebellar cell types^1^. Pilocytic astrocytomas, even though representing benign tumors, are often associated with life-long morbidities and multiple tumor recurrences^17^. While the origin of these tumors is unknown, PAs are hypothesized to arise from oligodendrocytes^18^. Notably, the cellular origins of pediatric tumors have been primarily deduced from murine cell atlases^1,2^, a limited and potentially imperfect approach due to spatio-temporal developmental gene expression differences between these species^3,19^. Therefore, it is essential to revisit the cellular origins of pediatric tumors based on a comprehensive human cell atlas.

To fill this gap, we compared transcriptomes representing cell types, differentiation states, and subtypes from the developing human cerebellum (*Sepp, Leiss, et al*.) to bulk and single-cell gene expression profiles from pediatric CNS tumors (**Figure 1a, Supplementary Figure 1a**). Our integrated analyses of these datasets revealed the best matching cellular lineages of origin for medulloblastoma, ependymoma and pilocytic astrocytoma classes and subclasses. Using tumor-to-normal tissue comparisons, we identified candidate target genes and pathways as putative tumor-specific vulnerabilities. Additionally, we detected the glial lineage, a source of both astrocytes and oligodendrocytes, as the lineage of origin in radiation-induced gliomas^20^, a relatively common secondary tumor arising after medulloblastoma treatment^21^, suggesting that these tumors arise independently rather than occuring through evolution or trans-differentiation from the primary medulloblastoma. To enable readers to access and use our resources and results further, we developed an online graphical user interface, which facilitates the interactive exploration of our data and results at *brain-match.org*.

**Figure 1.**
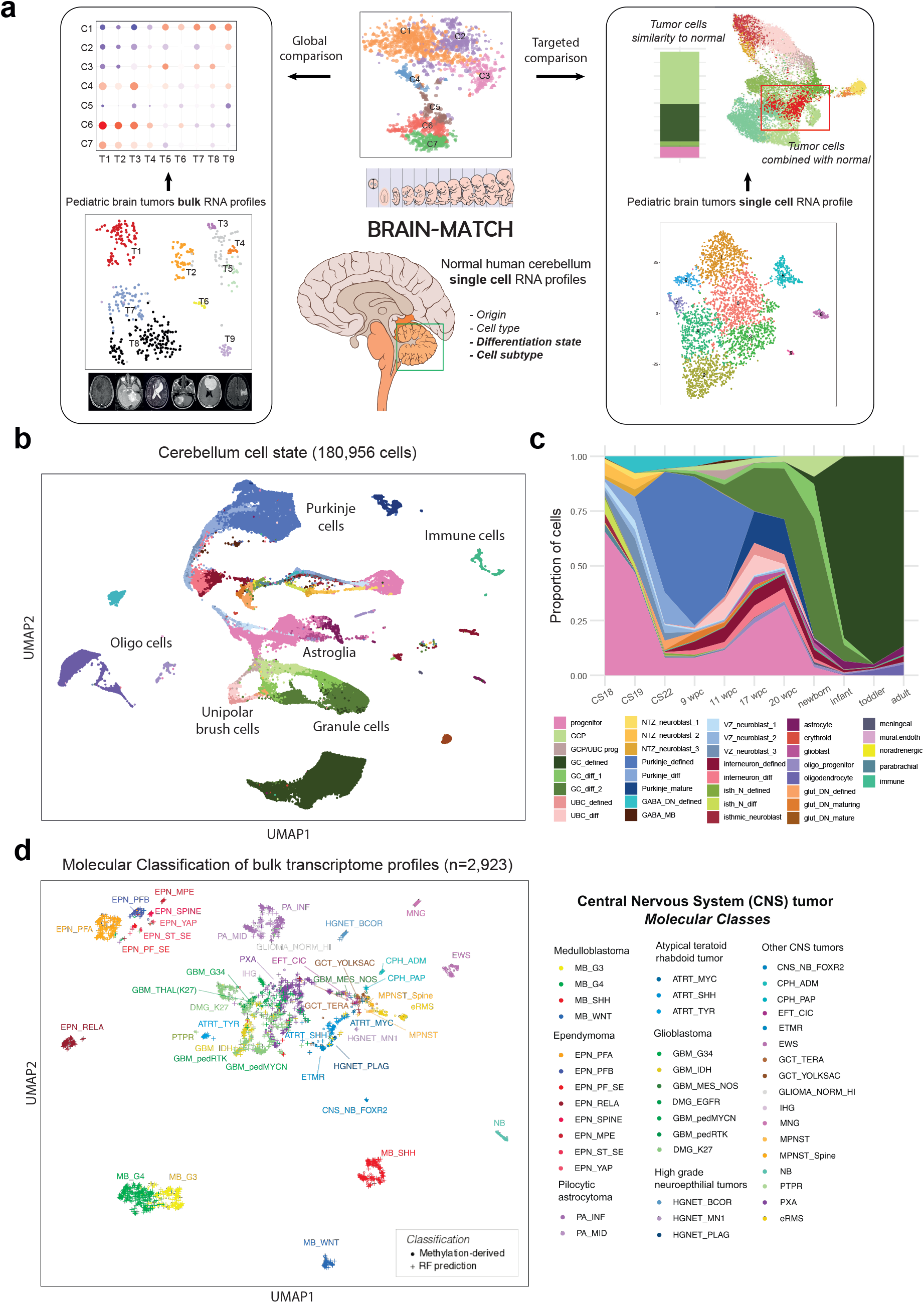
Human single-nuclei developmental cerebellum cell atlas as the reference for deciphering lineages-of-origin of cerebellar tumors. a) Overview of the BRAIN-MATCH project. Single-nuclei gene expression profiles of the developing brain region(s) serve as the reference for comparing to region-specific bulk or single-cell tumor gene expression data. b) Uniform Manifold Approximation and Projection (UMAP) visualization of human developmental cerebellum single-nuclei data. Color coded by the cell state annotation and labels show the main cell types. c) Proportions of different cells in different cell states across time points of collected data. d) UMAP visualization of main bulk tumor cohort colored as per methylation classification. Random forest based predicted samples depicted in cross.

## Distinct origins for different tumor classes

In the companion paper (*Sepp, Leiss, et al*), we generated an extensive single-nucleus RNA sequencing (snRNA-seq) dataset that covers the development of the human cerebellum from the beginning of neurogenesis to adulthood, and serves as a reference for the tumor-matching analyses in this study. In the reference atlas, 180,956 cells (**Figure 1b**) are grouped into 21 cell types (e.g. astroglia), further divided into 37 cell differentiation states (e.g., progenitor), hereafter referred to as cell states (**Suppl. Figure 1a**; **Figure 1c**). In addition, subtypes are specified for cell types and states, resulting in 65 transcriptomically-defined subtypes of cells (e.g., progenitors divided in nine spatio-temporal subtypes including rhombic lip progenitors). Notably, cell states within a cell type form a continuum along the differentiation trajectory, which we refer to as “cellular lineage” in this study. For instance, the astroglial lineage includes progenitor (neuroepithelial and radial glial cells), glioblast, and astrocyte cell states.

We used an unprecedented set of Affymetrix bulk expression data as our main tumor transcriptome cohort (**Suppl. Figure 1b)**. To remove sample-or study-specific classification biases, we used methylation-based classification, covering 55% of the target samples; for samples with missing methylation data, we used a random forest model to impute missing methylation classes (**Suppl. Figure 1c-d**). This Affymetrix data cohort represents 2,923 samples annotated into 45 defined molecular classes (e.g. medulloblastoma G3) and 68 methylation subclasses (e.g. medulloblastoma G34_II) (**Figure 1d, Suppl. Table 1**). We supplemented this Affymetrix cohort with an independent collection of bulk RNA-seq datasets (**Suppl. Table 2**), the largest of them the from the Children’s Brain Tumor Network (CBTN, n = 626) (**Suppl. Figure 1e**).

After considering existing computational approaches^1,2,22-24^, we selected two main measures to determine the similarity of a tumor class to a normal cell type: correlation of expression patterns of shared highly-variable genes^23^ and gene set variance analysis (GSVA)^24^. These measures were validated using SHH medulloblastoma as a positive control, given that this tumor class was experimentally shown to arise from granule cell progenitors (GCPs)^13,14^. A direct comparison of our training medulloblastoma RNA-seq dataset^25^ (ICGC cohort, n = 160) to cerebellar cell types also confirmed these results (**Suppl. Figure 2a**). Additionally, we found that at least 400 cells are needed per reference cluster to obtain a stable score when using our combinatorial correlation and GSVA approach (**Suppl. Figure 2b-c**), and we estimated the filtering thresholds for these computed values (**Suppl. Figure 2d-e**).

We next compared the large Affymetrix transcriptome tumor cohort to cerebellar cells (**Figure 2**). We included non-cerebellar tumor classes, such as meningioma (MNG), or glioma profiles enriched with immune cells (GLIOMA_NORM_HI) as external controls for this global comparison, which demonstrated matches to the expected cell states.

**Figure 2.**
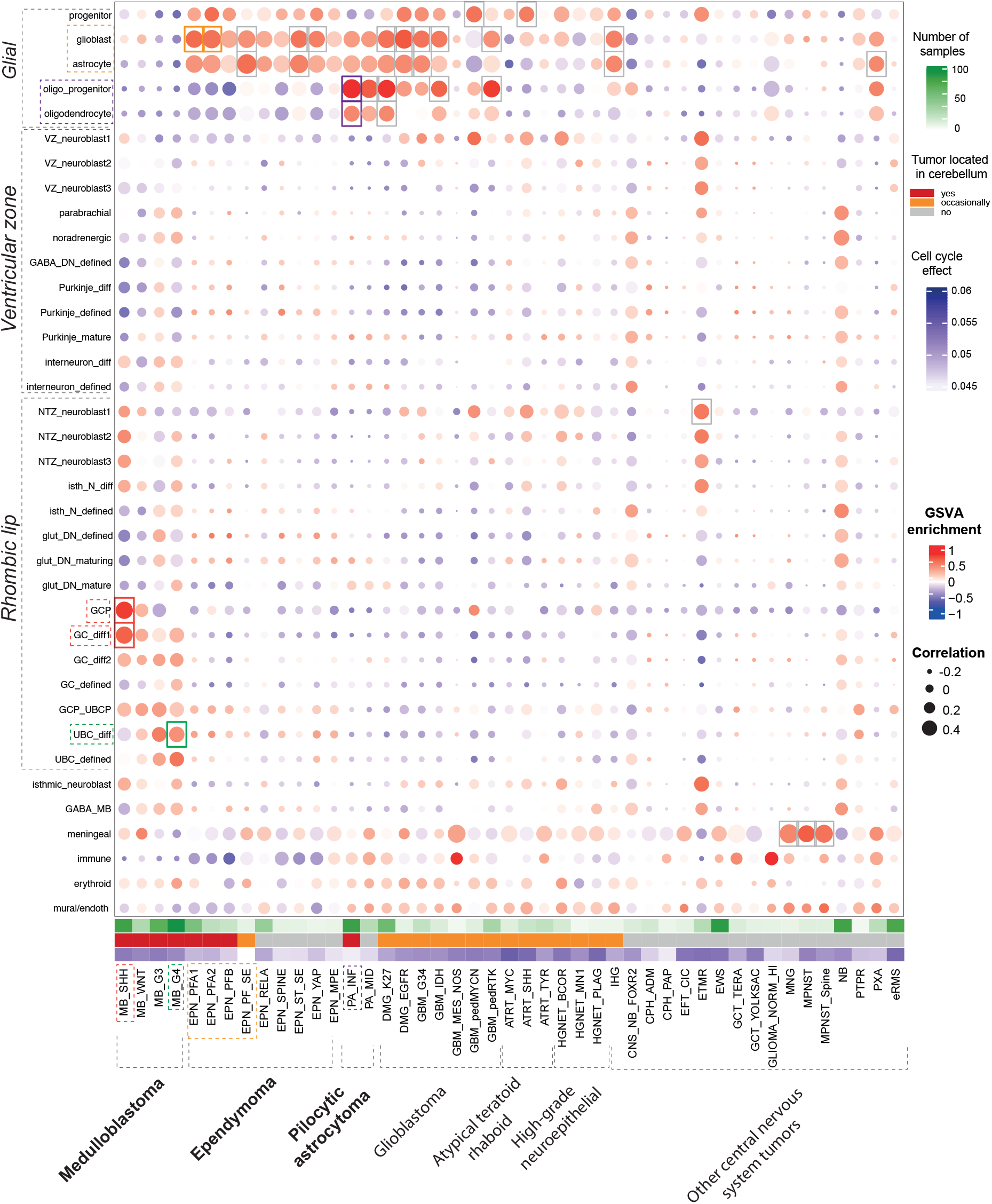
Global analysis identifies lineages-of-origin for multiple tumor types. Heatmap showing comparisons of bulk central nervous system tumor transcriptome profiles (in columns) to cerebellar *cell state* (in rows), based on gene signature enrichment score (via Gene Set Variance Analysis (GSVA), indicated by color) and Pearson correlation score (indicated by area). Most evident association of a tumor type to a cell state is highlighted by dashed rectangles, selected based on cutoff limits (min GSVA enrichment = 0.4 and min correlation = 0.4).

From this comparison, we further confirmed the closest association of SHH MB to GCP and differentiating granule cells (GC_diff1). Group 4 MB most closely matched differentiating unipolar brush cells (UBC_diff), confirming previous studies using murine atlases for comparison^1,2^. Associations of Group 3 medulloblastoma were ambiguous and showed several matches within the glutamatergic lineage, including progenitors and differentiating cells of the GC/UBC lineage, amongst others, but none of them passed our filtering thresholds. Predictably, WNT-medulloblastoma did not match to any cell state of the developing cerebellum, as it is believed to arise from the developing brainstem^12^.

The two classes of posterior fossa A (PFA) ependymoma, PFA1 and PFA2, were most similar to cells in the astroglial lineage, which includes progenitors, glioblasts and astrocytes. We note that some glial cell types including ependymal were missing in our human cerebellum dataset. Other posterior fossa ependymoma tumor classes, which are much rarer in children, including posterior fossa B (PFB) and posterior fossa subependymoma (PF_SE), showed similar associations with the astroglial lineage.

Infratentorial pilocytic astrocytoma (PA_INF), a tumor class mostly occurring in the cerebellum, showed the closest association with oligodendrocyte cell states – oligodendrocyte progenitor cells (OPC) and oligodendrocytes, thus validating a previous hypothesis about their cellular origin^18^. Several glioblastoma (GBM) molecular classes only occasionally located in the cerebellum^26,27^, demonstrated similarities to different glial cell states, such as astrocytes, oligodendrocytes, and their respective precursors.

Additional comparisons of tumors at the methylation subclass level recapitulated the detected associations at the molecular class level; for example, MB SHH subgroups were clearly related to GCP (**Suppl. Figure 3a**). The results we obtained using the CBTN cohort as a validation dataset were fully compatible with what we had found using the Affymetrix data (**Suppl. Figure 3b**), thus validating our approach using different expression datasets and platforms.

After obtaining these global results, we next sought to investigate the cellular origins of pediatric tumors in greater detail and at single-cell resolution focusing on the three most common entities arising in the cerebellum.

## Posterior fossa ependymomas likely arise from the astroglial lineage

PFA and PFB, the two major classes of ependymoma located in the cerebellum/posterior fossa were previously proposed to originate from radial glial cells^16^. This result was partly confirmed in a murine cerebellum-based reference map that found an association with “roof-plate like stem cells” and “gliogenic progenitors”^1^, but it has not been validated previously against a human reference. Using our human cerebellum single-cell dataset, we found that the transcriptional profiles of posterior fossa ependymomas most closely resembled various states of differentiation within the astroglial lineage of the cerebellum starting from radial glial and neuroepithelial progenitors and ranging to glioblasts followed by astrocytes (**Figure 2, Suppl. Figure 4a**), which in turn showed similarity to the “roof-plate-like stem cells” and “gliogenic progenitors”, as defined in a published murine cerebellum atlas^1^ (**Suppl. Figure 4b**), thus potentially resolving the apparent contradiction.

To increase the resolution of the cell type associations, we next focused on cell subtypes within the astroglial lineage (**Figure 3a**). There are two spatially segregated gliogenic paths in the cerebellum that start from gliogenic progenitors (producing Bergmann glia and parenchymal astrocytes) and bipotent progenitors (producing GABAergic interneurons and parenchymal astrocytes), respectively, and progress via their corresponding glioblast subtypes towards mature astrocytes^28^ (*Sepp, Leiss, et al*). We found that transcriptomes matching bipotent progenitors, prospective white matter glioblasts (glioblast_PWM), astroblast, and mature parenchymal astrocyte subtypes signatures were enriched in both PFA and PFB ependymoma samples (**Figure 3b**).

**Figure 3.**
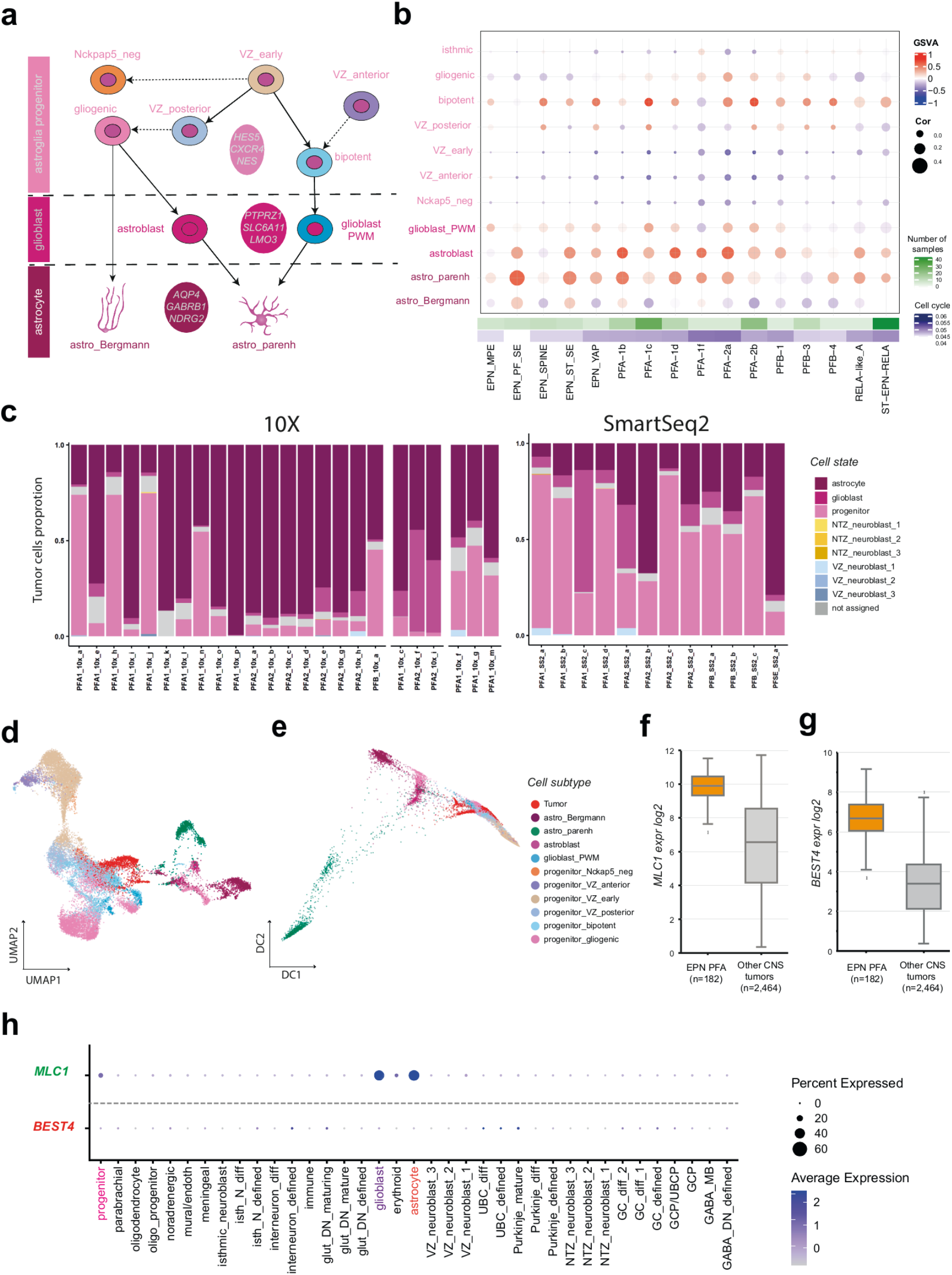
Ependymoma tumors correspond to the astroglial lineage. a) Genealogy of astroglial cell lineage subtypes as derived from early ventricular zone (VZ) progenitors. b) Comparison of bulk ependymoma tumor gene expression profiles to cell subtypes of the astroglial lineage, based on GSVA enrichment and correlation measures. c) Ependymoma tumor single cell data SingleR-SVM comparison to cerebellum cell state per sample. d,e) Example showing integration of single cell tumor data (ID: PFA1_10x_g, 10x v2) with astroglial cell subtypes as visualized via UMAP (d) and DiffusionMap (e) f) Expression boxplot of ependymoma PFA specific gene shared with astroglial lineage *MLC1* (*limma* adjusted p-value: 5.67E-31). The center line, box limits, whiskers and crosses indicate the median, upper/lower quartiles, 1.5× interquartile range and outliers, respectively. g) Expression boxplot of tumor-unique ependymoma PFA specific gene *BEST4* (*limma* adjusted p-value: 1.75E-61). The center line, box limits, whiskers and crosses indicate the median, upper/lower quartiles, 1.5× interquartile range and outliers, respectively. h) Examples of genes shared between ependymoma PFA and astroglial lineage (green), and tumor-specific (red), gene expression across the cerebellum cell states

Based on these observations, we hypothesize that either the tumor develops from the earliest node within the astroglial lineage and differentiates along the specified trajectory after malignant transformation, or that the transcriptome of posterior fossa ependymoma cells is comprised of a mix of progenitor and mature cell signatures. To investigate these possibilities further, we investigated published posterior fossa ependymoma single-cell data from three independent studies^1,29,30^. We assigned each tumor cell to its closest matching cell state identity, using an adjusted SingleR-SVM approach^31^ (see Methods for details, **Suppl. Figure 4c**). After assignment, we found non-malignant immune cell clusters in PF ependymoma samples analyzed using 10x chemistry (**Suppl. Figure 4d**). As expected, these cell clusters also had normal DNA copy number profiles when compared to tumor cell clusters (**Suppl. Figure 4e**); hence, we excluded them from further analysis to prevent obscuring the tumor signature.

Similarly, we also filtered out cells with assigned identities that did not align to the astroglial lineage (for example, meningeal, GCP, oligo-progenitors, etc.), given that these tumor cells do not form independent cell clusters and therefore may be rare non-immune normal cells from the tumor microenvironment. After filtering, we found that each tumor sample had cells resembling progenitor, glioblast, and mature astrocyte-like cells, but with different proportional distributions within tumors of the same class (**Figure 3c**). This result suggests that a tumor is comprised of cells along a differentiation trajectory within the lineage of origin. We tested this hypothesis by overlaying tumor cells onto the astroglial lineage and found that the tumor cells laid along the entire lineage, suggesting a gradient of differentiation within PF tumors with a possible origin from bipotent and gliogenic progenitors (**Figure 3d, e**). Surprisingly, the transcriptomic signatures of progenitor-like and astrocyte-like tumor cells did not show mutually exclusive marker gene expression as seen in the normal lineage stages; however, enrichment of marker genes (e.g. progenitor: *TENM3*, astrocyte: *AQP4*) supported the assigned identity of these tumor cells (**Suppl. Figure 4f**). We thus predict that even “mature” tumor cells maintain proliferative capacity. We also observed a similar gradient of differentiation in an independent Smart-seq2 dataset (**Suppl. Figure 4g**). Altogether, and with the caveat of not having ependymal cells represented in our dataset, our findings suggest that PF tumors are composed of cells along a maturation gradient of the astroglial lineage.

We next determined shared genes between the cerebellar astroglial lineage and PFAs (**Supplementary Table 3**). Among the shared genes, we focused on those that are specifically expressed in the brain and/or cerebellum^3^, and either confirmed to be plasma membrane localized or possible drug targets based on their Human Protein Atlas annotation^32^. As one of the top candidates, we identified the potentially druggable *MLC1* gene expressed in ependymomas (**Figure 3f**), which was also expressed specifically in the astroglial lineage, but not in any other cell lineage in the cerebellum (**Figure 3f)** or outside of the brain (**Suppl. Figure 4h**). Similarly, we identified and filtered tumor-specific candidates including the gene *BEST4*, which encodes a membrane protein and was not detected in any cerebellar cell lineage (**Supplementary Table 4, Figure 3g,h**); it is also absent or lowly expressed in other normal tissues (**Suppl. Figure 4i**). These two potential therapeutic targets could be explored for CAR-T therapies to specifically attack tumor cells^33^. Through further investigation of affected pathways, we observed specificity in neurogenesis for genes common between tumors and the astroglial lineage, while, cancer-related pathways (e.g. GBM silenced by methylation or soft tissue tumors activity) were enriched in tumor-specific gene sets (**Suppl. Figure 4j**).

## Pilocytic astrocytomas correspond to postnatal OPC

Our global comparisons confirmed previous observations that two pilocytic astrocytoma molecular classes, PA_INF (infratentorial or posterior fossa, predominantly cerebellum) and PA_MID (supratentorial midline), most closely resembled oligodendrocyte progenitor cells (OPCs) (**Figure 2**).

We examined an additional bulk RNA-seq cohort^34^ covering another class of pediatric low-grade gliomas, namely supratentorial gangliogliomas (GG_ST), which also most closely resembled OPCs (**Suppl. Figure 5a**). We observed from our global comparisons that diffuse midline gliomas, H3 K27 altered (DMG_K27) were associated with astrocytes and also with oligodendrocytes (**Figure 2**); therefore, we included DMG_K27 tumors as an external comparison for further investigation of PA origins. We then examined the subtypes along the oligodendrocyte lineage and found the best match to late oligodendrocyte progenitor cells (OPC_late) that are present in the postnatal cerebellum (**Figure 4a,b; Suppl. Figure 5b**). The similarity level demonstrated an association with the anatomic location of the tumor in the brain, i.e. posterior fossa PA showed the closest match to cerebellar OPCs, with a less striking match in midline PAs. Using single-cell gene expression data from independent cohorts^1,18^ (non-tumor cells excluded as described above), we confirmed the similarity of transcriptomes between pilocytic astrocytoma and OPCs (**Figure 4c**). We then overlaid single-cell tumor data onto the oligodendrocyte lineage and found that most tumor cells integrated with the OPC_late cluster, with some cells overlapping with early OPCs and mature oligodendrocytes (**Figure 4d,e; Suppl. Figure 5c)**.

**Figure 4.**
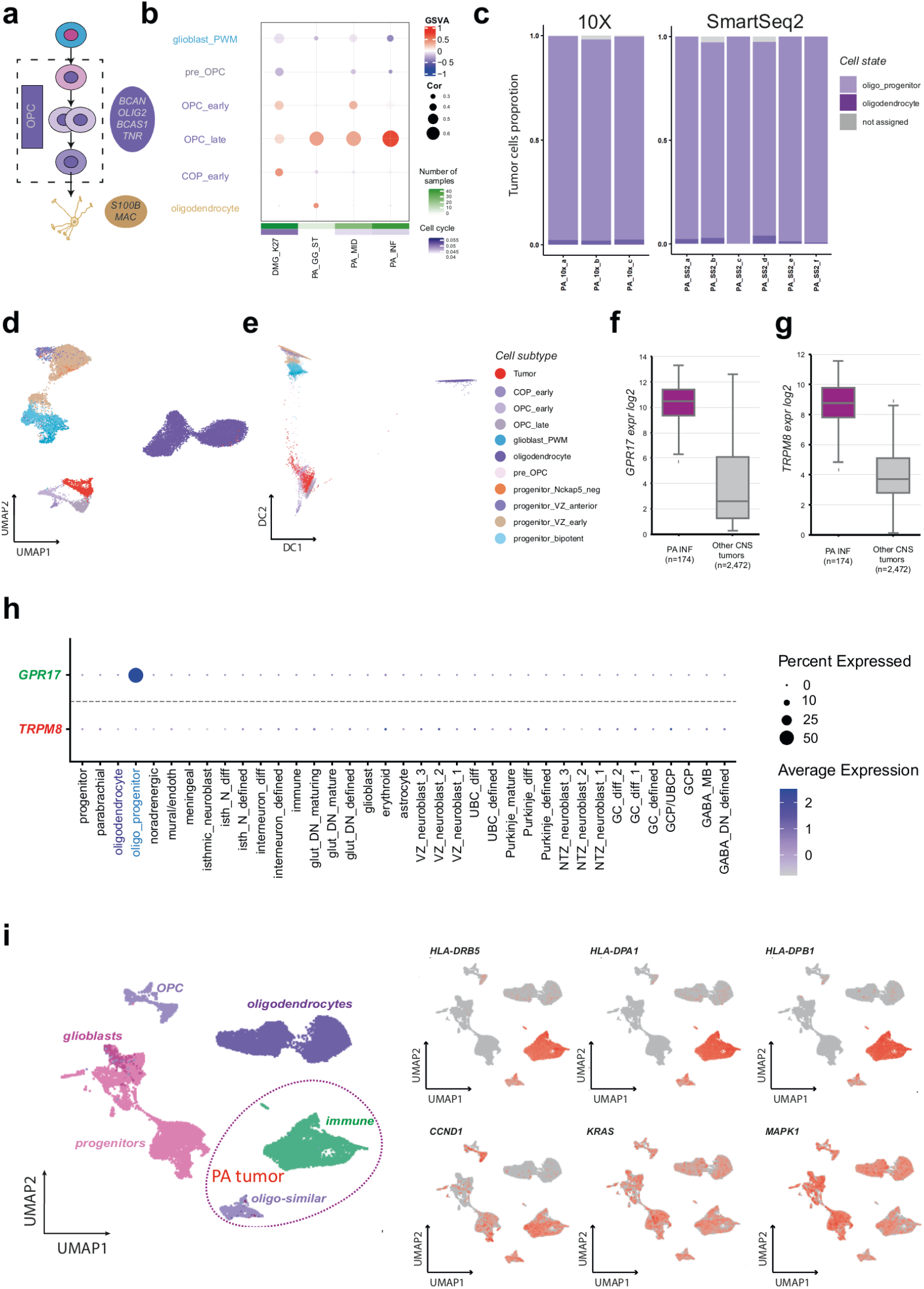
Pilocytic astrocytomas arise from the oligodendrocyte lineage. a) Genealogy of oligodendrocyte cell lineage subtypes starting from bipotent progenitors b) Comparison of bulk PA tumor gene expression profiles to cell subtypes from oligodendrocytic lineage based on GSVA enrichment and correlation measures. K27M DMG is external control. c) PA tumor single cell data correlation-based comparison to cerebellum cell state per sample. d,e) Example showing integration of single-cell tumor sample (ID: PA_10X_a, 10x v2) with oligodendrocyte cell subtypes as visualized via UMAP (d) and DiffusionMap (e). f) Expression boxplot of PA specific gene shared with oligodendorcytic lineage *GPR17* (*limma* adjusted p-value: 1.25E-105). The center line, box limits, whiskers and crosses indicate the median, upper/lower quartiles, 1.5× interquartile range and outliers, respectively. g) Expression boxplot of tumor-unique PA specific gene *TRPM8* (*limma* adjusted p-value: 2.14E-72). The center line, box limits, whiskers and crosses indicate the median, upper/lower quartiles, 1.5× interquartile range and outliers, respectively. h) Examples of genes common in PA and oligodendrocyte lineage (green) and tumor-specific (red), gene expression across cerebellum cell states i) Direct combination of single-cell sample (ID: PA_10X_a, non-tumorous immune cells included) with oligodendrocyte lineage

**Figure 5.**
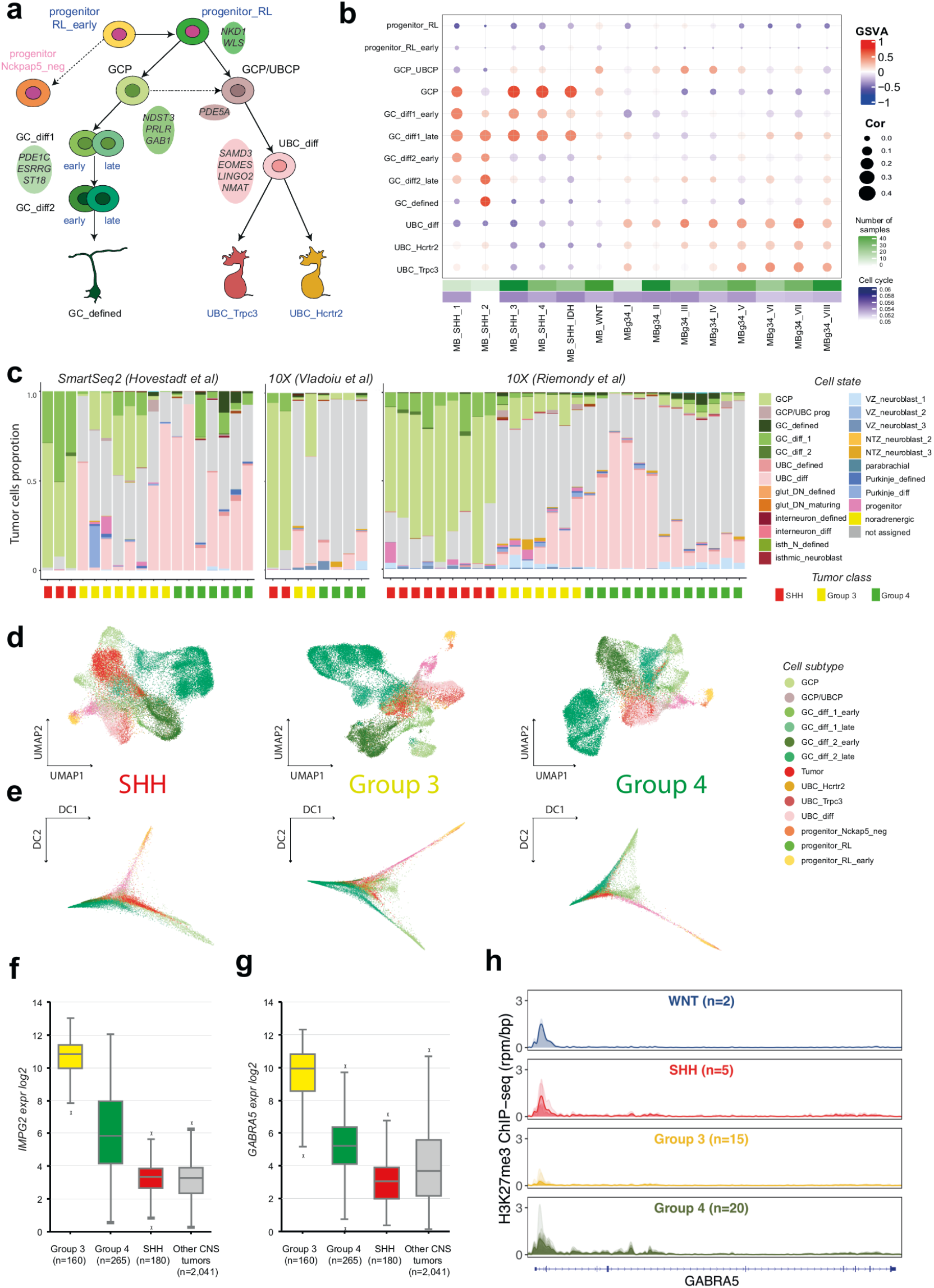
Medulloblastomas correspond to granule/unipolar brush cell lineages. a) Genealogy of granule/unipolar brush cell (GC/UBC) lineage subtypes derived from rhombic lip progenitors. b) Comparison of bulk medulloblastoma gene expression profiles to the cell subtypes from GC/UBC cells, based on GSVA enrichment and correlation measures. c) Medulloblastoma single cell data correlation-based comparison to cerebellum cell state per sample. d,e) Examples of 10X tumor samples (IDs: SHH_10X_b, G3_10x_d, G4_10x_f) showing integration with the cell subtypes in granule/unipolar brush cell lineage as visualized via NMF UMAP (d) and DiffusionMap (e). f) Expression boxplot of tumor-unique medulloblastoma specific gene *IMPG2* (*limma* adjusted p-value: 5.72E-148). The center line, box limits, whiskers and crosses indicate the median, upper/lower quartiles, 1.5× interquartile range and outliers, respectively. g) Expression boxplot of tumor-unique medulloblastoma specific gene *GABRA5* (*limma* adjusted p-value: 2.89E-77). The center line, box limits, whiskers and crosses indicate the median, upper/lower quartiles, 1.5× interquartile range and outliers, respectively.h) H3K27me3 ChIP-seq profiles (reads per million mapped reads per bp, rpm/bp) of *GABRA5* loci in the four MB classes.

We next assessed shared genes between pilocytic astrocytoma and OPCs (**Suppl. Table 3**), and identified some notable candidates, such as *GPR17*, which encodes a cell-surface protein (**Figure 4f,h; Suppl. Figure 5d**). Inspection of tumor-unique genes also revealed some interesting candidates (**Suppl. Table 4**), including potentially druggable *TRPM8* as a possible candidate for therapeutic intervention (**Figure 4g,h; Suppl. Figure 5e)**.

Using gene-set enrichment analysis, we found that OPC and pilocytic astrocytomas shared genes involved in oligodendrocyte development, while interestingly tumor-specific genes were specifically enriched in major histocompability complex class II (MHC II) and immune system associated genes (**Suppl. Figure 5f**). Because this immune signature could result from immune cells within the tumor micro-environment, we inspected single-cell data from pilocytic astrocytomas. After excluding non-malignant cells, we still found expression of MHC class II genes (e.g., *CD74, HLA-DRB5*) in tumor cells (**Suppl. Figure 5g**), similar to previous observations in pilocytic astrocytoma for some MHC class I members^18^; thus these genes are indeed overexpressed in tumor cells. Notably, known downstream targets of MAPK signaling (pathway constitutively activated in pilocytic astrocytoma), including *CCND1* (Cyclin D), *KRAS* or *MAPK1* were expressed across most normal and tumor cell clusters and within the oligodendrocyte lineage (**Figure 4i**). The immunological role of this pattern in terms of tumorigenesis and possibly contribution to the senescence phenotype typically observed in PA remains to be studied.

## Medulloblastomas originate from the GC/UBC lineage

From our global comparison, SHH, Group 3, and Group 4 MBs closely associated with one or both branches of the GC/UBC lineage (**Figure 2**); therefore, we next focused on a high-resolution lineage map to delineate the potential origins of different medulloblastoma subclasses (further referred also as subgroups based on WHO terminology^6^) (**Figure 5a**). We found that most of the SHH-medulloblastoma subgroups closely corresponded to GCPs and nascent postmitotic GCs (GC_diff1) (**Figure 5b**). SHH-medulloblastoma subgroups I,II,IV showed high similarity to the late subpopulation of differentiating GCs that emerge at postnatal stages (GC_diff1_late). SHH-II (beta) subgroup, which is associated with infant onset and metastasis^35^, instead resembled more differentiated GC cells states including defined GC (GC_defined) that have reached the inner granule cell layer.

Group 3 and 4 medulloblastoma subgroups range from I-VIII, with subgroups II/III restricted to Group 3 and subgroups VI/VIII restricted to Group 4^36^. Subgroups V-VIII (mostly Group 4 medulloblastoma) best matched to UBCs (differentiating and defined). Subgroups I-IV (mostly Group 3 medulloblastoma) displayed low transcriptome correlations with subtypes in the GC/UBC lineage; nevertheless, GSVA enrichment scores suggested some similarity with GC/UBC progenitors and early differentiating UBC. The lowest correlation/GSVA similarity to normal cell types was observed for Group 3/4 subgroup II, which is known to be associated with *MYC* amplification^36^. Such somatic *MYC* changes may drive the tumor to further deviate from the original lineage as investigated in more details below.

Using medulloblastoma single-cell data^1,2,37^, we recapitulated these findings (**Figure 5c**). Group 4 tumor cells aligned with differentiating GCs and UBCs, suggesting that these tumors differentiate along a bifurcating GC/UBC trajectory, with a predominance of UBC-like cells (**Figure 5d,e; Suppl. Figure 6a)**. Similarly, we found that Group 3 tumor cells differentiated along a similar trajectory, but with a predominance of proliferating and differentiating GC-like cells. Remarkably, both Group 3 and Group 4 medulloblastoma samples had a high proportion of unassigned cells (less than 50% classification probability and classified as “not assigned”), but were most similar to GC/UBC-like cells based on best-matching identity (**Suppl. Figure 6b**). Consistent with these observations, recent animal studies proposed the transforming capacity of *Atoh1*-positive lineages such as GCs and UBCs into *MYC*-driven Group 3^38^ and *SRC*-driven Group 4 tumors^39^.

**Figure 6.**
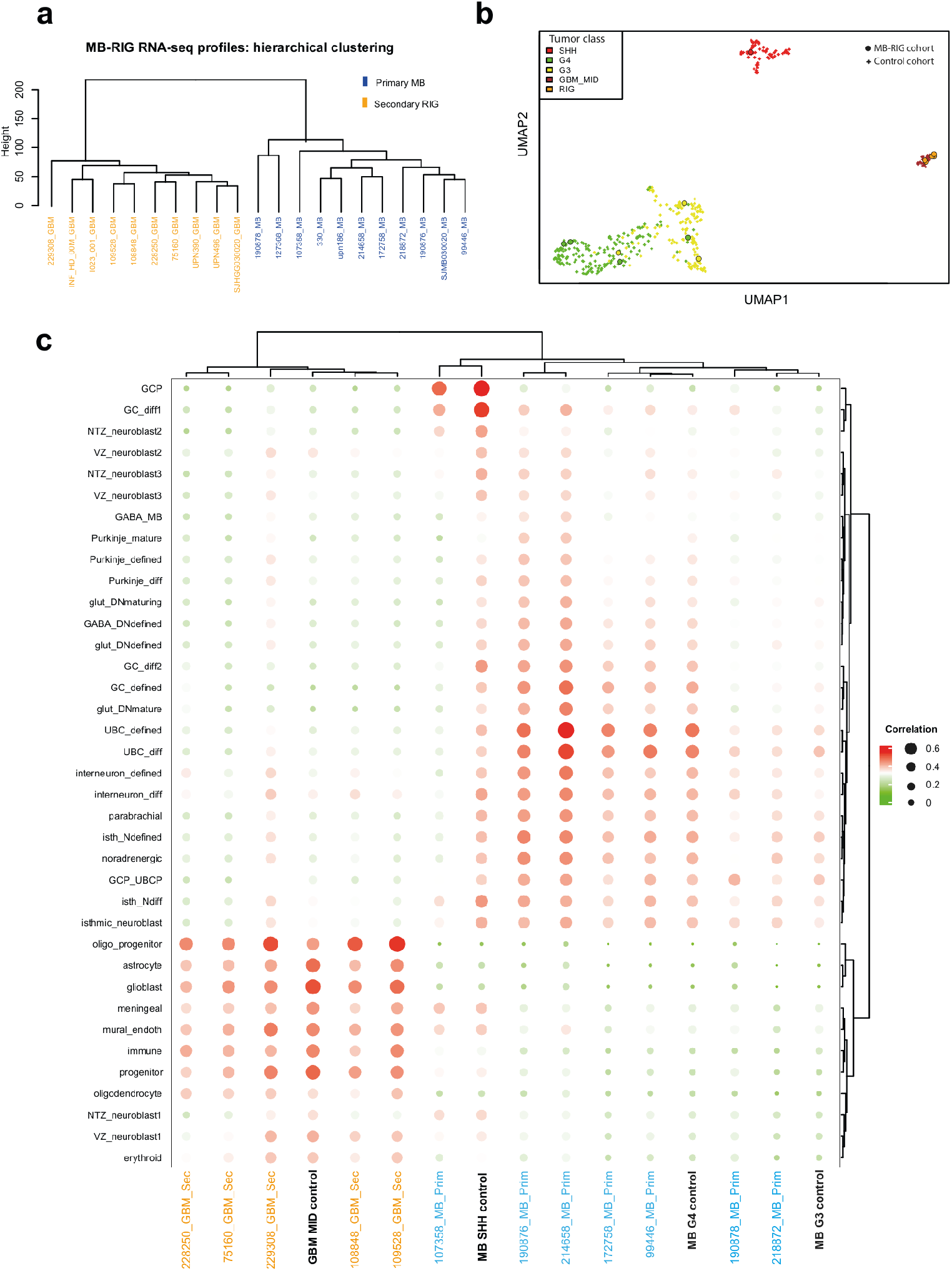
Secondary radiation induced gliomas (RIGs) originate from the glial cell lineage and not from the original primary tumor. a) Unsupervised hierarchal clustering of batch-effect adjusted RNA-seq profiles from primary medulloblastoma and secondary RIG tumors (based on top 500 HVGs) b) UMAP visualization of medulloblastoma and glioblastomas. Primary medulloblastoma and corresponding secondary RIGs highlighted with circles. c) Comparison of gene expression profiles of primary medulloblastoma and secondary RIG tumors combined with corresponding FFPE bulk control to cerebellum cell state, based on correlation measures.

Some Group 3/4 tumor cells best matched to differentiating Purkinje cells or nuclear transitory zone neuroblasts, suggesting the presence of cells outside of the GC/UBC lineages. However, these tumor cells did not cluster separately from other GC/UBC-like cells and expressed GC/UBC lineage specific genes (**Suppl. Figure 6c**,**d**), suggesting the expression of multi-lineage markers leading to this ambiguous identity. Alternatively, somatic alterations in proliferative genes driving tumor development, such as *MYC/MYCN* amplifications (like in Group 3 subgroup II as described above), could possibly explain this phenomenon, and may prevent the tumor cells from resembling a defined lineage. A similar phenomenon has been shown in mice^38^. In addition, when separately examining *MYC* (Group3) and *MYCN* (SHH and Group 4) amplified tumors, they did not show the expected similarity to the GC/UBC signature, confirming the impact of these somatic changes (**Suppl. Figure 6e**).

To identify clinically relevant candidates, we focused on genes shared between the medulloblastoma classes and the GC/UBC lineage (**Supplementary Table 3**). An example of shared genes, *EOMES*, a transcription factor specific for Group 4 medulloblastoma, is known to play an important role in brain development and has strong association with structural defects in the fetus^40^ (**Suppl. Figure 7a-c**). Among possible tumor unique candidates (**Supplementary Table 4)**, we identified the surface protein coding gene^37^, *IMPG2*, which is specific for Group 3 tumors and lowly expressed in normal tissue (**Figure 5f; Suppl. Figure 7a,d**).

We next identified the gene ontology pathways detected in these gene groups to further inspect their molecular associations (**Suppl. Figure 7e-g)**. Even though association with CNS development was found across all medulloblastoma classes, Group 3 tumor-specific genes were highly enriched for H3K27me3-suppressed genes in normal stem cells (**Suppl. Figure 7f**). Some of these genes e.g., *GABRA5* (**Figure 5g**), were found to be present only in tumors. Changes in H3K27me3 have been observed to play a role in Group 3 medulloblastoma formation^41,42^; notably, we also found loss of H3K27me3 in these genes specifically in Group 3 (**Figure 5h, Suppl. Figure 7h,i**).

## Radiation-induced gliomas are independent secondary tumors based on lineage of origin

Radiation-induced gliomas (RIGs) may occur as a consequence of cranial radiotherapy commonly used to treat CNS tumors^43^, including medulloblastoma^44^. Based on their glioblastoma-specific genetic landscape, the lack of overlapping mutations with the initial medulloblastoma^21,45^, and on functional studies in non-human primates^46^, it is assumed that these tumors arise *de novo* from healthy cells after radiation rather than by trans-differentiation of residual medulloblastoma cells. However, this has never been formally demonstrated. To further investigate this hypothesis, we focused on radiation-induced gliomas (n = 11) and the respective primary medulloblastoma samples (n = 11) for comparison to the cerebellum cell types (**Suppl. Table 5, Suppl. Figure 8a**). As expected, copy number variation analysis showed distinct, non-overlapping profiles for primary and secondary tumors, with an increased number of insertion and deletion events in RIGs (**Suppl. Figure 8b**). Unsupervised hierarchical clustering of gene expression profiles further distinguished primary medulloblastomas from secondary RIGs (**Figure 6a**). For a subset of formalin-fixed paraffin-embedded (FFPE) primary-relapse matched samples, we compared transcriptome profiles with an additional FFPE bulk RNA-seq dataset (n = 410), representing sporadic medulloblastoma and glioblastoma samples (**Suppl. Table 2**). UMAP visualization of the tumor pairs with control data resulted in grouping of the primary and secondary tumors with medulloblastoma and GBM clusters, respectively (**Figure 6b**).

We next compared each medulloblastoma-RIG pair to cerebellar cell states, using correlation analyses only (GSVA enrichment requires differentially expressed genes from >2 samples), including merged profiles from sporadic medulloblastoma and glioblastoma datasets. RIG tumors, along with glioblastoma controls, correlated with glioblast/astrocyte/oligodendrocyte cells comprising the glial lineage, whereas medulloblastoma primary tumors and controls were most similar to the cells in the GC/UBC lineage (**Figure 6c**). We also merged all RIG samples into a single group and computed both correlation and GSVA scores; these samples corresponded to oligodendrocyte progenitors and glioblasts. From this data, we conclude that the lineage-of-origin is different between primary medulloblastoma and secondary RIG tumors (**Suppl. Figure 8c**), and that secondary RIGs arise exclusively from a glial lineage.

## Discussion

Using an unprecedented cell atlas of the developing human cerebellum, we performed a comprehensive comparison of childhood brain tumor cohorts to normal cells and determined the most probable cellular lineages of origin for the most common tumor types arising in the cerebellum. One of the remarkable observations from our exhaustive analysis was that rarely any tumor class exhibited associations with only one cell state. Instead, tumors typically contained a gradient of differentiating cells along a cellular lineage. However, tumors belonging to a class did show compositional heterogeneity in terms of similarity to normal cells. In addition, alignment of tumors to the proposed lineage of origin showed differences in the starting point of tumorigenesis along a normal cell type lineage trajectory. These findings suggest that tumors from the same class may arise at several differential stages in the associated lineage, and that tumor expansion often follows a pre-specified trajectory, which closely resembles the developmental trajectory of the lineage of origin. Importantly, it also indicates that tumors exploit developmental and functional characteristics of the lineage of origin to grow and develop into their complex form, similar to a tissue containing cells of different functions.

For posterior fossa ependymomas, we found a possible origin in the astroglial lineage, covering previous studies using murine cerebellar atlas^1^, in which however only “roof-plate-like stem cells” (possibly choroid plexus or ependymal precursors) and gliogenic progenitors were hypothesized as the cell-of-origin. We also found transcriptomic similarities between these murine cell types to a subset of progenitors in the human cerebellum atlas. Nevertheless, in the future, expanding to regions outside of the cerebellum, including posterior hindbrain regions, would help to further refine this observation and uncover cell types that were missed, such as ependymal cells, potentially also in connection to ependymomas^29^.

In our analysis, pilocytic astrocytomas associated with postnatal OPCs. The favorable prognosis of these patients could be a result of slow growth, MAPK-induced senescence, increased immunogenicity, or a combination thereof. Importantly, we also found that MHC class II members are abundantly expressed in tumor cells, which suggests these tumors may be better recognized by the immune system, in contrast to other CNS tumors.

For SHH-, Group 3-, and Group 4-medulloblastomas, we found possible origins within the two branches of the GC/UBC lineage. While these tumors are all associated with the same broad lineage, the relative proportions of GC and UBC cells within a tumor varied depending on the tumor class/subclass. We found that while Group 4 tumors are enriched with UBCs, these tumors also harbor the potential to differentiate along the GC trajectory. This finding suggests that Group 4 tumors arise from an earlier precursor cell that is able to generate both UBC and GC. Group 3 medulloblastomas were also composed of both GC and UBC cells; however, they were not enriched in either signature. Instead, many cells within Group 3 tumors were ‘not-assigned’ after filtering, or exhibited a non-GC/UBC lineage signature (e.g., NTZ neuroblasts or differentiating Purkinje cells) along with GC/UBC marker genes. The inability to assign a specific identity to Group 3 tumor cells could occur if the cell type of origin was not captured in our cerebellar atlas or it deviates a lot from all normal lineages based on strong MYC activity. We also observed that Group 3 was associated with specific H3K27me3 loss and strong activation of a group of genes that are typically silenced in normal stem cells. One example, *GABRA5*, has a clear connection with aggressive tumor growth^47^ and appears to be ubiquitously expressed within the tumor, making it a potentially attractive target for therapeutic interference.

We identified the most specific marker genes that are expressed in tumors and the closest lineage of origin, as well as genes that are tumor-specific and are not detected in the developing cerebellum. The main reason for this combined inspection was that even though tumor-active genes not present in normal tissues were initially suggested to serve as ideal targets, some recent studies challenged this view^48^, while the use of genes that are active not only in the tumor, but also in progenitor cells demonstrated better results (e.g., *CD133* in gliomas^49^). In addition, some lineage-specific genes remain active only in progenitors within a lineage (e.g., *GPR17*, active in OPCs only), and hence could be used as candidate targets based on their persisting expression in the resulting tumors. Finally, a lineage-specific gene active in mature cells could be used for gating strategies to target localized tumors^50^. Importantly, if these tumor-specific genes or lineage-shared genes are not detected or are only minimally expressed in postnatal non-brain tissue, and if the gene expresses a protein localizes to the plasma membrane, this target could be used in next generation CAR-T therapies.

Current limitations in the understanding of oncogenic vulnerabilities of normal cell types and tumor progression therefrom, are major roadblocks in search of new specific therapies in pediatric neuro-oncology. In our work, we focused on uncovering these conundrums by integrating a human single cell cerebellum atlas to pediatric CNS tumors via global comparisons. We have made the analysis tools and results freely available via an interactive graphical interface at *brain-match.org* for further hypothesis generation and exploitation of these precious resources for the benefit of our patients.

## Supplementary Figures

**Suppl. Figure 1.**
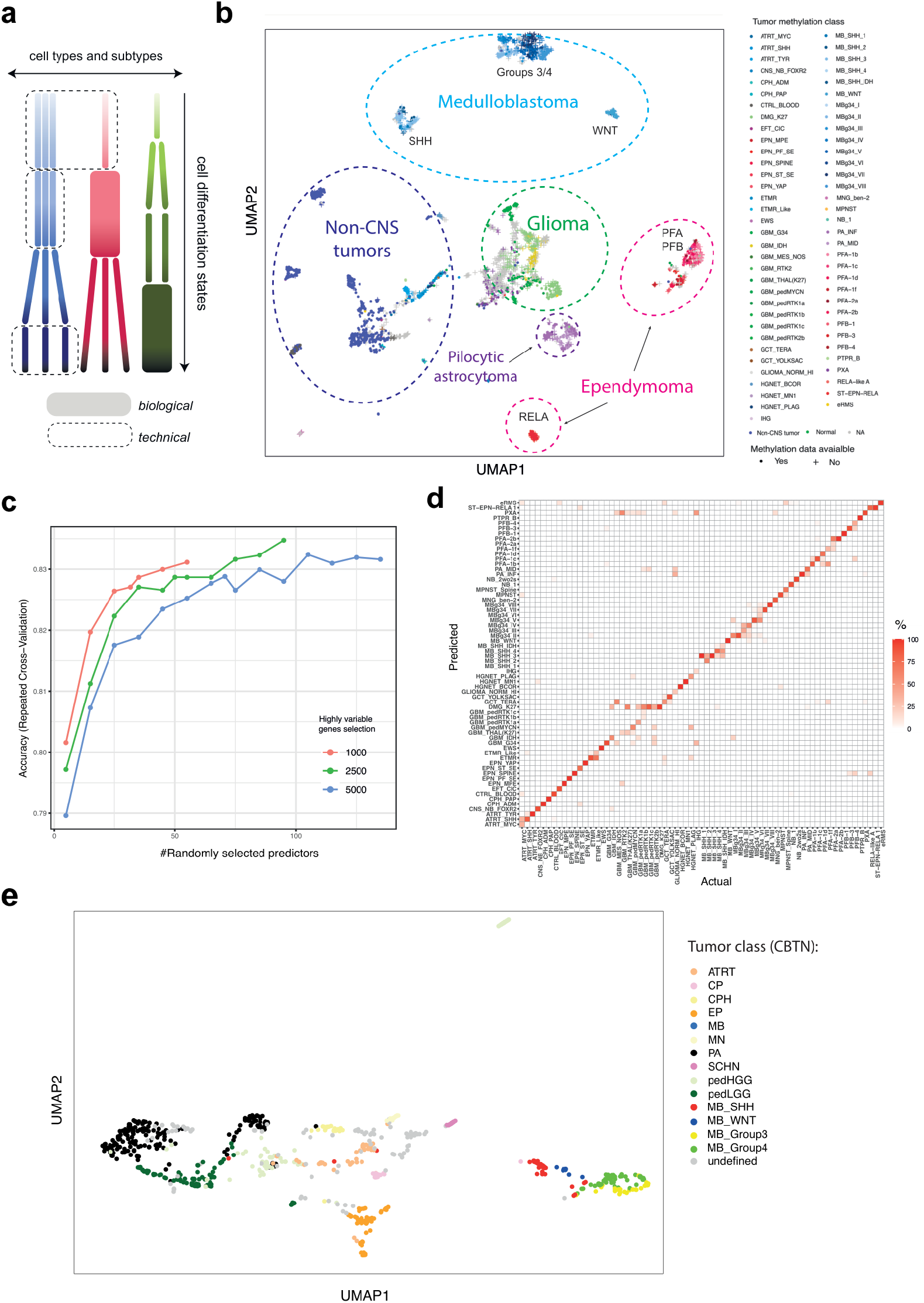
Global combined profiles of cerebellar tumors and their classification. a) Schematic summary of the strategy used for the annotation of cells in the reference atlas of human cerebellum development. Shown are 3 hypothetical cell types (reflected by color) and their subtypes (rectangles). The y-axis depicts differentiation that gives rise to a continuum of cell differentiation states. Biological and technical reasons (legend below the plot) could explain cases when subtypes cannot be distinguished across all states in a given cell type. b) UMAP representation of combined Affymetrix tumor cohort (based on top 500 HVG) color coded as per methylation classification mnp12. Random forest predicted samples are marked by cross. c) Statistical details of random forest classification showing dependency of accuracy on number of predictors d) Heatmap showing results of a 5-fold cross-validation of the random forest classifier incorporating information tumor samples with methylation data. e) UMAP representation of CBTN pediatric brain tumor RNA-seq dataset (based on top 2000 HVG).

**Suppl. Figure 2.**
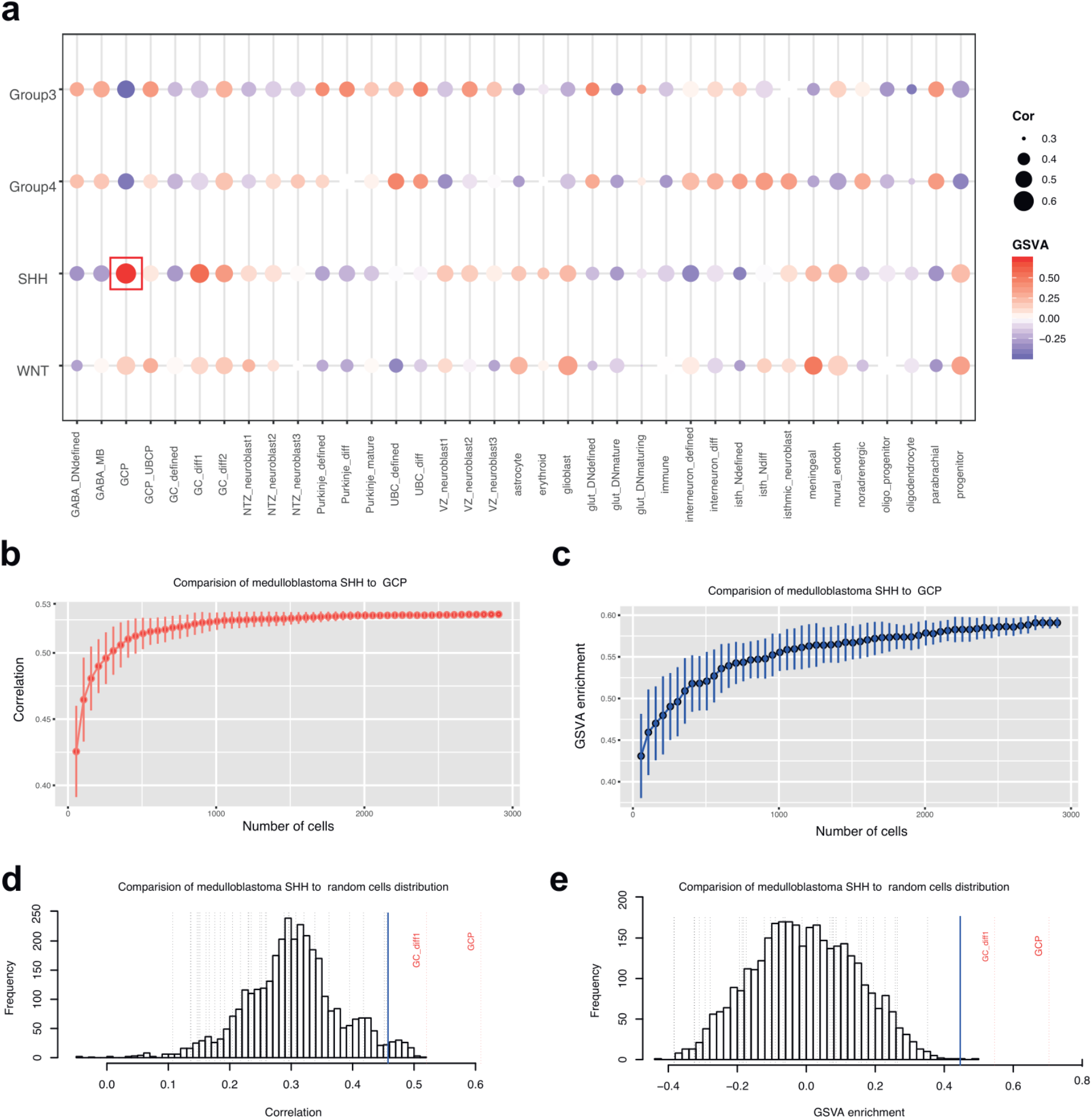
Validation of GSVA and correlation measures strategy to find correspondence of tumor class with closes cell state match. a) Comparison of ICGC MB RNA-seq data to cerebellum cell states based GSVA enrichment and correlation values. Size of circle: correlation measure; Color of circle: GSVA enrichment measure. b-c) Effect of cluster size on correlation (b) and GSVA(c) measures stability: size of GCP cluster was varied from 100 to 3000 cells randomly selected cells, showing correspondence to SHH medulloblastoma plateaued around ∼300-500 cells. d-e) Cutoff limits for correlation (d) and GSVA(e) measures: clusters derived from permutated cells (100 repeats, random cluster size reflects sizes of cell state classification) are compared to SHH medulloblastoma bulk RNA-seq data. Dashed lines represent result achieved from cell state comparisons. Cutoff limit is derived from empirical rule adjustment (mean + 3 standard deviations). Mean cutoff limits computed from all MB classes comparison; correlation: ∼0.4, GSVA enrichment: ∼0.4

**Suppl. Figure 3.**
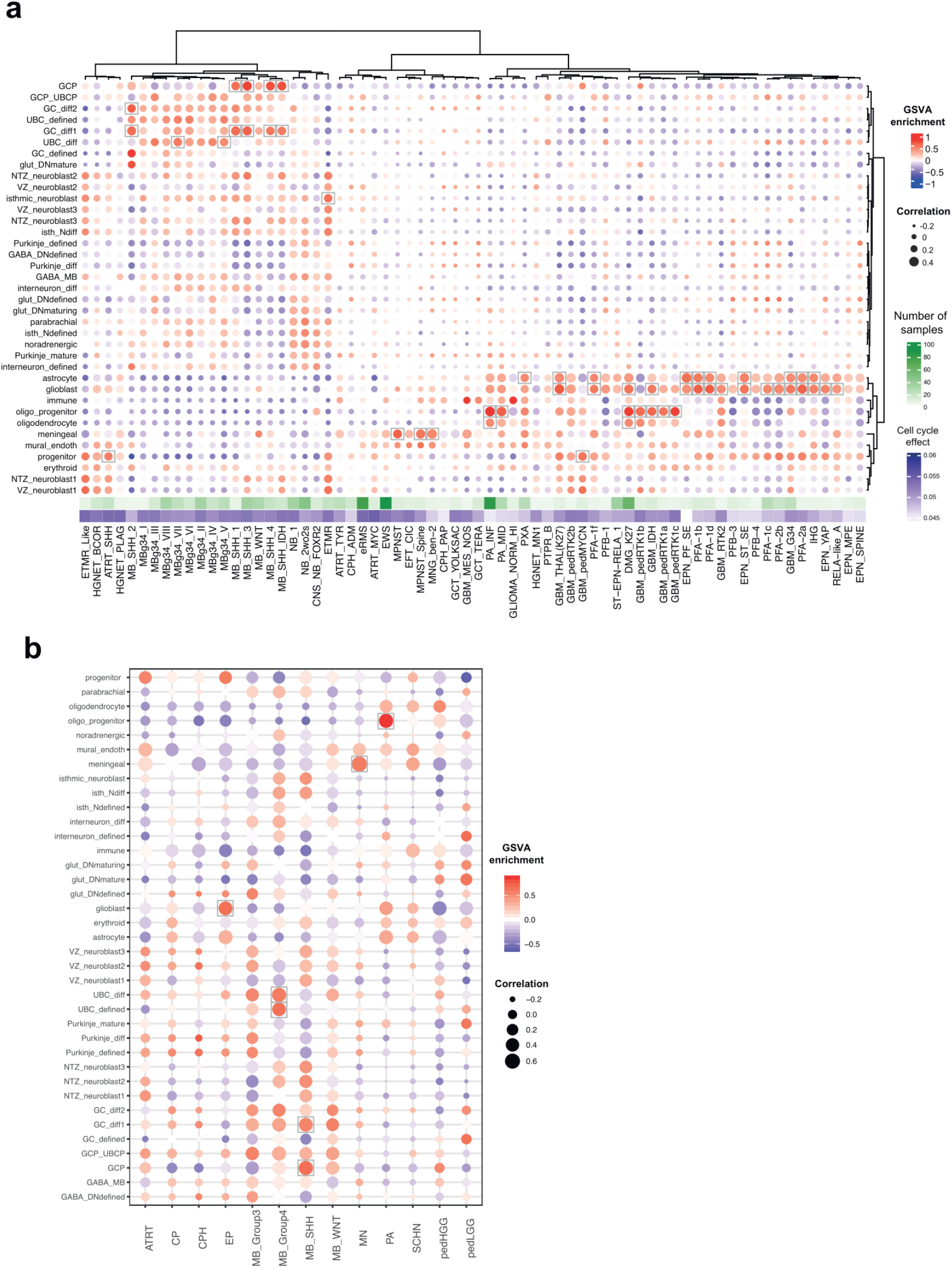
Extended global tumor data comparisons confirm identified lineages-of-origin for cerebellum-specific CNS tumors. a) Comparison of main tumor bulk gene expression profiles in Methylation Classes (mnp12.3) to normal cerebellum cell states based on GSVA and correlation measures. Unsupervised clustering based on correlation measure is applied to group tumor classes and cell types. b) Comparison of CBTN RNA-seq gene expression profiles to normal cerebellum cell states based on GSVA and correlation measures, recapitulating tumor-normal developmental state associations obtained from Affymetrix data. The observations obtained from mnp12.3 global cohort comparison are fully confirmed.

**Suppl. Figure 4.**
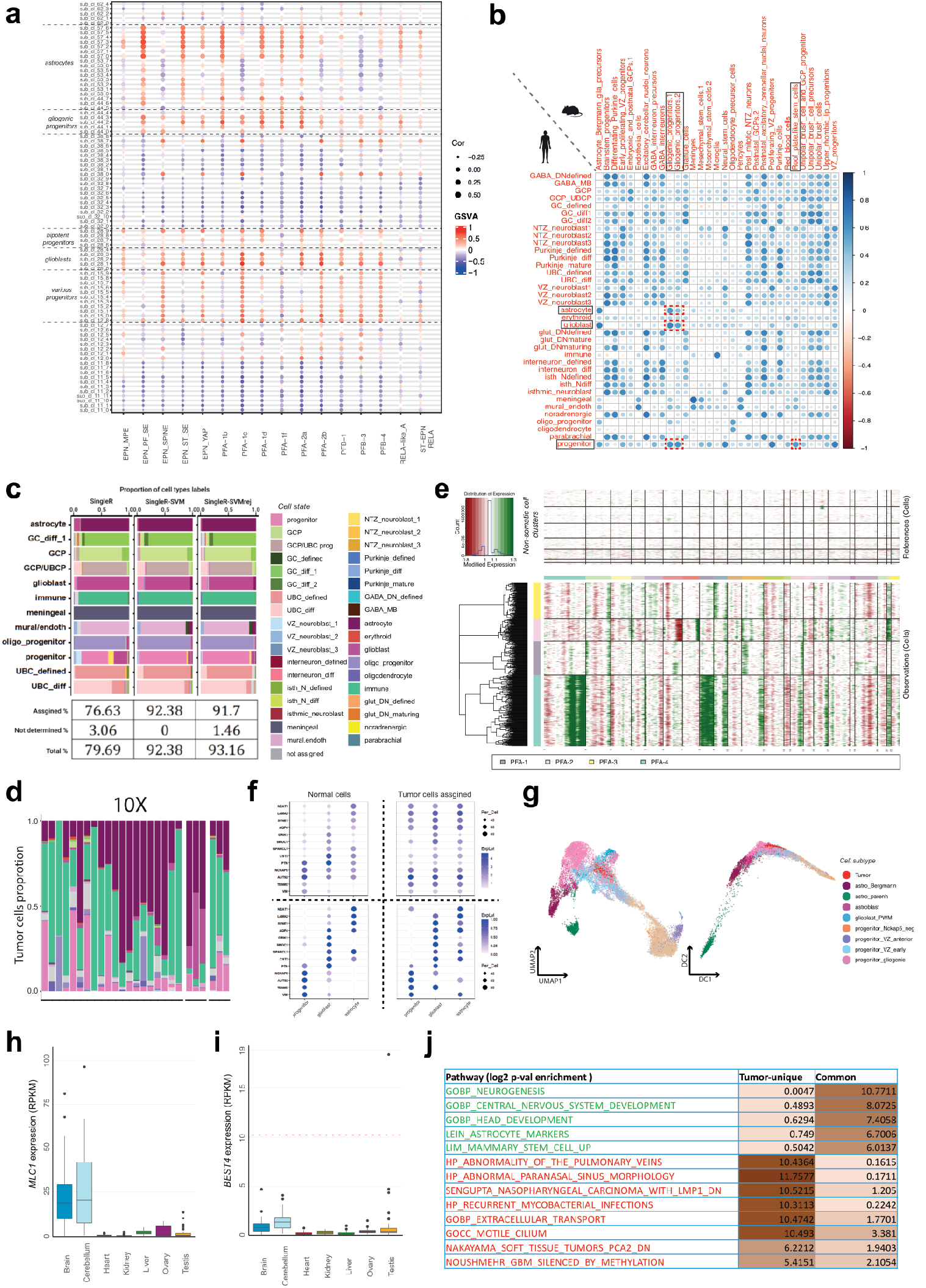
Ependymoma tumors correspond to the astroglial lineage. a) Comparison of gene expression profiles of ependymoma tumor classes to CB cell subclusters from astrocyte lineage based GSVA enrichment and correlation measures. b) Correlation-based comparison of mouse cerebellum cells vs human cells based on usage of orthologous genes. c) Verification of SingleR-SVM based label assignment of cell types using train (80%) and test (20%) sets of cerebellum data. SingleR: SingleR based correlation measure; SVM: SingleR assignment pruned with SVM; SVMrej; SVM based best matches filtered for 50% probability cutoff. d) Ependymoma single cell dataset (10x) comparison to cerebellum per sample via SingleR-SVM. Enrichment of immune cells observed across samples. e) CNV profiles of ependymoma 10X samples allow to distinguish somatic and non-tumor cells f) Mutually exclusive expression pattern of marker genes in normal progenitor, glioblast and mature astrocyte (left side), that are co-expressed in progenitor-like, glioblast-like and astrocyte-like tumor cells (right side). Scaling (below), however, show why based on pattern of expression the cell identity assigned is comparative to normal cell types. g) Integration of PFA1 sample (PFA1_SS2_a) with astroglial lineage shown using UMAP and DiffusionMap. h) Expression boxplot of ependymoma PFA specific gene *MLC1* across normal tissues. The center line, box limits, whiskers and crosses indicate the median, upper/lower quartiles, 1.5× interquartile range and outliers, respectively. i) Expression boxplot of ependymoma PFA specific gene *BEST4* across normal tissues. The center line, box limits, whiskers and crosses indicate the median, upper/lower quartiles, 1.5× interquartile range and outliers, respectively. j) Top enriched GO pathways for ependymoma PFA specific DEGs either shared with astroglial lineage or uniquely enriched in PFAs.

**Suppl. Figure 5.**
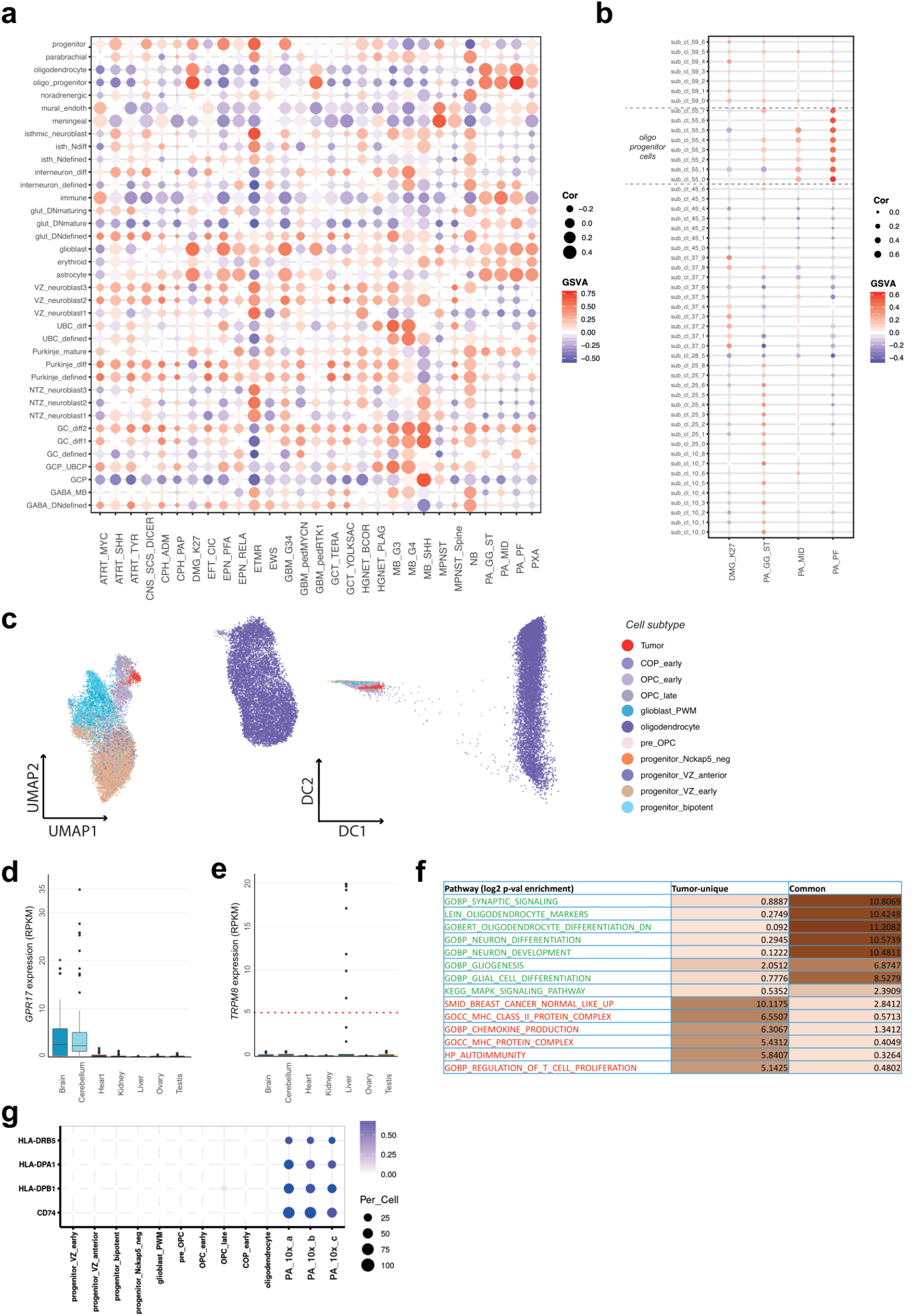
Pilocytic astrocytomas arise from the oligodendrocytic lineage. a) Comparison of INFORM RNA-seq profiles to normal cerebellum cell states based on GSVA and correlation measures. b) Comparison of gene expression profiles of PA tumor classes to CB cell subclusters from oligodendoryte lineage based GSVA enrichment and correlation measures. c) Integration of single cell PA data (PA_SS2_a) VZ progenitors and oligodendrocyte lineage as shown using UMAP and DiffusionMap d) Expression boxplot of PA specific gene *GPR171* across normal tissues. The center line, box limits, whiskers and crosses indicate the median, upper/lower quartiles, 1.5× interquartile range and outliers, respectively. e) Expression boxplot of PA specific gene *TRPM8* across normal tissues. The center line, box limits, whiskers and crosses indicate the median, upper/lower quartiles, 1.5× interquartile range and outliers, respectively. f) Top enriched GO pathways for PA_INF specific DEGs either shared with oligodendrocytic lineage or tumor specific. g) Expression of MHC class II members protein complex that are not active in cerebellum oligodenrocyte lineage subtypes, but present in PA tumor cells.

**Suppl. Figure 6.**
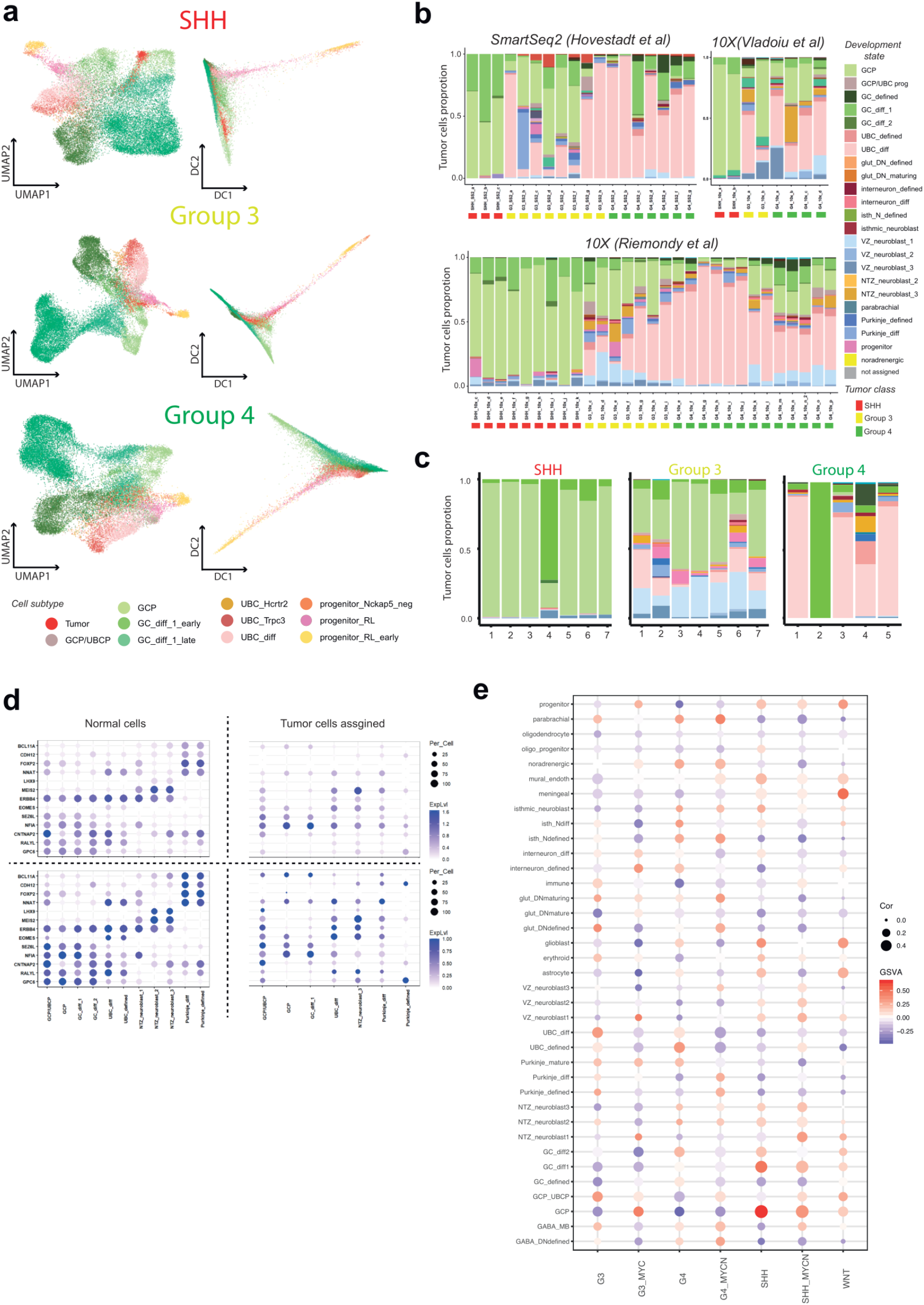
Medulloblastoma tumors correspond to granule/unipolar brush cell lineages. a) Integration of single cell SHH, Group 3 and Group 4 data with GC/UBC lineage as shown via UMAP and DiffusionMap. b) Medulloblastoma tumor single cell data in combination with main enrichment of granule cell and unipolar brush cells per sample also show presence of additional cell types (filtered non-assigned cell types for datasets MDT 10X, Gojo SS2, Riemondy 10X). c) Comparison of tumor cell clusters to cerebellum cell states d) Expression of NTZ_NB (*MEIS2, LHX9*) or Purkinje (*BCL11A*) expression is found in Group 3 tumor cells, giving them similar identity (right side). However, these cells also express GCP (*GPC6, NFIA*) or UBC (*EOMES*) markers, probably having a hybrid identity. e) Comparison of ICGC MB RNA-seq profiles with split by MYC (Group 3) and MYCN (SHH/Group 4) amplification status to normal cerebellum cell states based on GSVA and correlation measures.

**Suppl. Figure 7.**
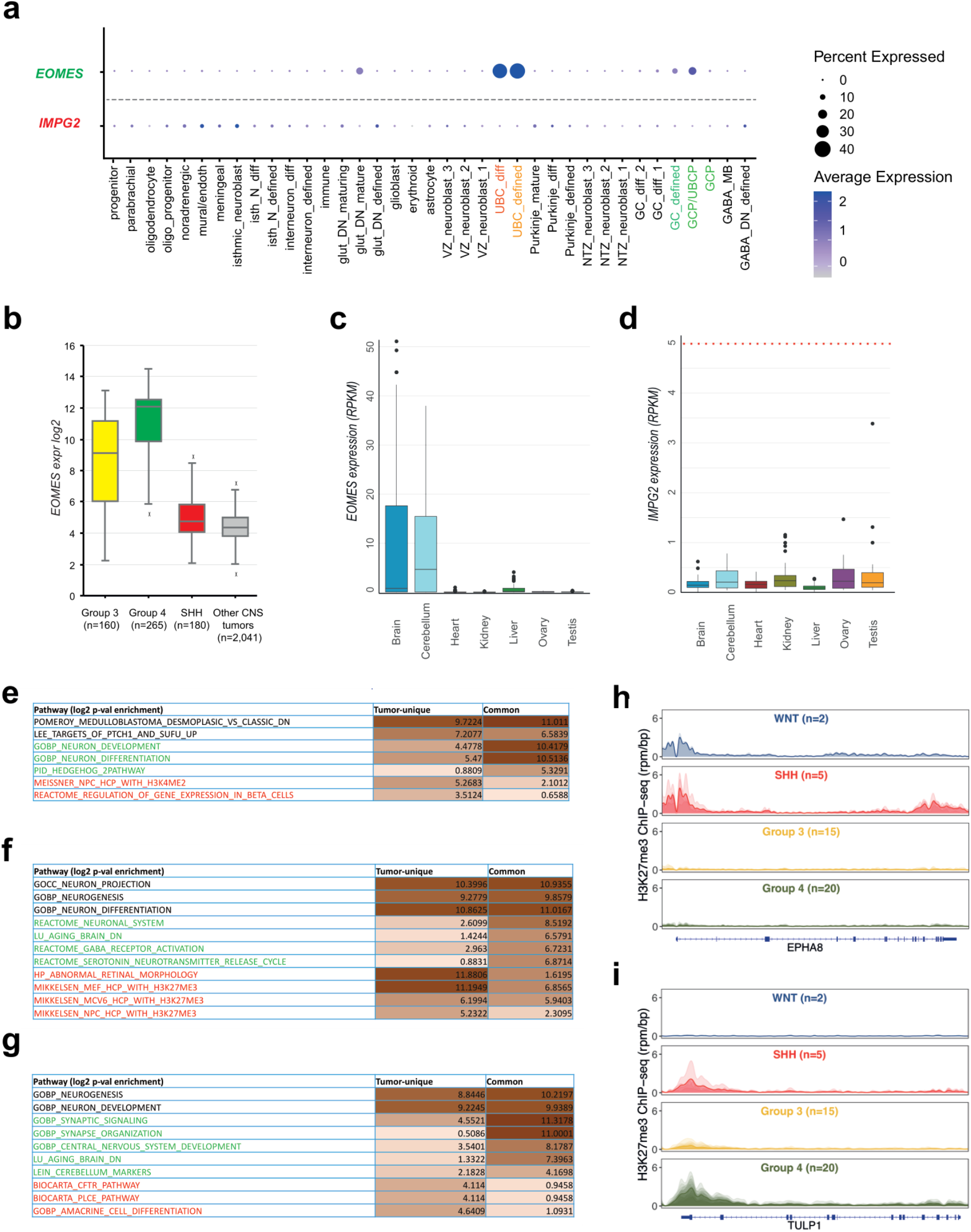
Medulloblastoma genes in correspondence to granule/unipolar brush cell lineages and unique tumor-specific. a) Examples of genes common for MB and GC/UBC lineage (green) and tumor-specific (red), gene expression across the cerebellum cell states b) Expression boxplot of medulloblastoma specific gene shared with GC/UBC lineage *EOMES (*differential expression adjusted p-value: 3.55E-27) c) Expression boxplot of MB G4 specific gene *EOMES* across normal tissues. The center line, box limits, whiskers and crosses indicate the median, upper/lower quartiles, 1.5× interquartile range and outliers, respectively. d) Expression boxplot of MB G3 specific gene *IMPG2* across normal tissues. The center line, box limits, whiskers and crosses indicate the median, upper/lower quartiles, 1.5× interquartile range and outliers, respectively. e) Top enriched GO pathways for SHH specific DEGs either shared with GC/UBC lineage or tumor specific. f) Top enriched GO pathways for Group 3 specific DEGs either shared with GC/UBC lineage or tumor specific. g) Top enriched GO pathways for Group 4 specific DEGs either shared with GC/UBC lineage or tumor specific. h,i) H3K27me3 ChIP-seq profiles (reads per million mapped reads per bp, rpm/bp) of EPHA8 (h) and TULP1 (i) loci in the four MB classes.

**Suppl. Figure 8.**
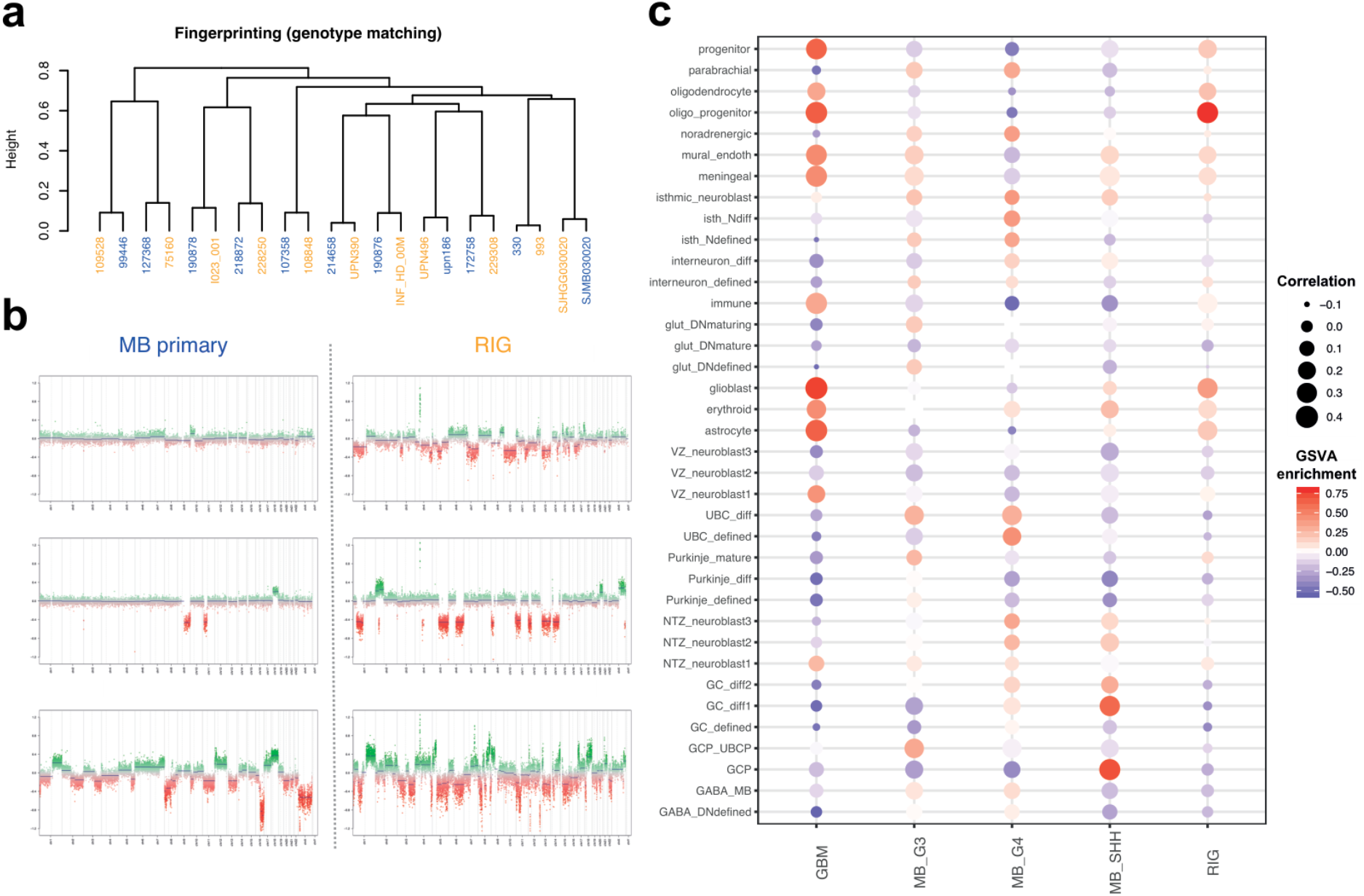
Secondary radiation induced gliomas (RIGs) originate from the glial cell lineage and not from the original tumor. a) Fingerprint match comparison for primary MB and secondary RIG tumors b) Copy number profiles of primary MB G3 vs secondary RIG tumors c) Comparison of FFPE RNA-seq gene expression profiles to normal cerebellum cell state cell types based on GSVA and correlation measures. Target primary secondary samples are included into MB cohort, while secondary RIG into GBM.

## Supplementary tables

**Suppl. Table 1**. Overview of the methylation and molecular mnp12.3 tumor classes.

**Suppl. Table 2**. List of bulk and single cell transcriptome tumor profiling datasets used in this study.

**Suppl. Table 3**. Differentially expressed genes of target tumor classes MB, EPN and PA common with genes specific for the cerebellum cell states.

**Suppl. Table 4**. Differentially expressed genes of target tumor classes MB, EPN and PA unique only for tumor and not/low expressed in the cerebellum.

**Suppl. Table 5**. Annotation of samples of primary MB and secondary RIG tumors

## Acknowledgments

This project has received funding from the European Research Council (ERC) under the European Union’s Horizon 2020 research and innovation programme (grant agreement no. 819894) and Seventh Framework Programme (FP7-2007-2013) (grant agreement no. 615253).

## Author Contributions

K.O. and P.J. performed bioinformatics analysis related to the comparison of tumors to normal cerebellum cell types. M.Se. and K.S. processed the data and provided detailed annotations for single nuclei human cerebellum materials. M.Se., K.S., I.S., F.M. and H.K. contributed to the project from evolutionary biology perspective. M.Si., P.B. and N.J provided additional tumor bulk data analysis materials. K.C.C., D.K. and M.Z. contributed to the gene selection control. A.K., F.S., M.Y.D., D.S., J.D. N.K.F., A.M.D., A.L.G., M.R., J.G., G.R., B.A.O., Q.G., E.D., J.L.H. provided tumor tissue samples along with processed data when possible and contributed to the design of the study. P.N., J.G., D.K., V.H., M.G.F., A.v.D., M.Z., K.W.P., M.K., D.T.W.J. and N.J. provided expert advice and helped to design the study. K.O., P.J., L.M.K. and S.M.P. prepared the figures and wrote the manuscript based on feedback from all authors. L.M.K, H.K., and S.M.P. conceived and co-led the study.

## Competing interests

The authors declare no competing financial interests.

## Methods

### Human cerebellum single nuclei RNA sequencing data

The snRNA-seq data generation and annotation are described in detail in the companion paper (*Sepp, Leiss et al*.). Shortly, 38 libraries from 31 independent human cerebellum samples were produced using Chromium (10x Genomics) version 2 or 3 reagents. The dataset includes 180,956 high quality cells from 10 developmental stages ranging from postconceptional week 7 to adult age. Louvain algorithm was used to cluster the merged dataset into 68 clusters and 611 subclusters. Hierarchical annotation of the dataset classified the cells into 21 cell types and 37 cell differentiation states. Cells from 12 cell states were further divided into 37 subtypes. For the purpose of the current study, we combined the cell state and subtype level annotations to divide the dataset into 65 subgroups. For simplicity, we refer to these subgroups as ‘subtypes’ throughout the study.

### Bulk tumor transcriptome data integration

The main Affymetrix tumor cohort used for our analyses was generated from the combination of 3 datasets: mixed CNS tumor cohort from various sources including normal brain (n=2351), glioblastoma tumor cohort (n=430) and relapse associated INFORM cohort with both CNS and non-CNS tumors (n=1040). After filtering for sample quality and exclusion of overlaps, the resulting cohort included 3497 unique samples.

Bulk RNA-seq profiles (Reads Per Kilobase of transcript, per Million mapped reads / RPKM, normalized gene expression counts) were obtained from four independent datasets: ICGC medulloblatoma^25^, Burdenko FFPE^51,52^, INFORM^34^ and Children’s Brain Tumor Network (*cbtn.org*).

For the Children’s Brain Tumor Network dataset gene expression counts were provided by the resource, while for remaining datasets main processing (alignment, gene expression counts computation, quality control) was performed as previously described^51^.

### Tumor profiles random forest classification and regression

In order to remove sample- or study-specific classification biases and assign DNA methylation-based classification mnp v12.3 (*molecularneuropathology.org/mnp*) to CNS tumor samples without this data type we performed a consensus classification of the entire tumor cohort. Samples covering both expression and methylation data (∼ 55% samples in the combined Affymetrix cohort) were used as a reference. Each reference mnp12.3 tumor class consisted of at least 5 samples (with minimum methylation-based assignment score >= 0.75 per sample), resulting in a total of 1199 samples across 68 tumor classes. We generated a random forest classifier based on the reference dataset with 5×5 cross validation via caret R package v6.0-86. Classification parameters including initial total number of genes, number of sample candidates (mtry) and number of trees were tested for precision and recall to select optimal settings. We then used the trained model to classify the remaining 1724 tumor samples lacking methylation data.

### Comparison of bulk tumor profiles to the normal cerebellum single nuclei data

We compared the transcriptomic signatures of tumor samples to those of normal cerebellum cell states or subtypes (as reference) based on a combination of correlation estimates and Gene Set Variance Analysis (GSVA^53^ package v1.28). Expression profiles of normal cerebellar cells were aggregated into pseudobulk across cell state and subtypes and normalized into RPKM values.

An intersection set of highly variable genes (HVGs) among tumor bulk profiles and cerebellum pseudobulk cell types computed via genefilter R package was used to calculate correlation. We optimized the number of HVGs used for the analysis using the well-established correspondence of SHH medulloblastoma to GNPs as a control, varying the number of HVGs from 1000 to 2000 with step 250 for cerebellum cell types and from 250 to 500 among tumor classes. For GSVA measure the differentially expressed genes among tumor classes within target bulk datasets were identified using limma^54^ package v3.42 with minimum adjusted p-value 0.05. The requirement was that tumor class of interest must have at least three samples for the detection of a specific differentially expressed gene signature^55^. Main control input variable for GSVA measures was number of selected differentially expressed genes specific for tumor class based on adjusted p-value < 0.05, and ranked by adjusted p-value for specificity. Score stability was observed between 50 to 200 genes, thus we used 100 DEGs as default for comparisons.

To inspect the effect of cluster size on correlation-GSVA measures, cerebellum single cell GCP cluster was subsampled with a minimum of 50 cells and a step size of 50 cells. Filtering cut limits for both measures were obtained from a distribution of computed correlation/GSVA enrichment scores obtained from random subsampling of cerebellum cells into possible clusters (repeated 100 times, number of cells selected based on proportions from annotations) in correspondence to SHH medulloblastoma and adjusted via empirical rule (*mean + 2*standard deviation*).

Cell cycle effect as annotation for bulk expression profiles was computed as proportion of expression of cell cycle genes^56^ in full expression profile across samples.

### Differential gene expression analysis among cerebellum cell types and tumor tissues

The differentially expressed genes among assigned cerebellum cell types were achieved via methods from Seurat v.3.2.2 (Wilcoxon Rank Sum and distance-based assignment). The genes differentially expressed among tumor classes were adjusted following selection of limma derived group-specific comparison (target tumor class vs all other samples in cohort) using minimum adjusted p-value 0.05 followed also with cut to top 2000 from the sorting. The tumor specific genes were compared to cell type via the overlaps by gene ID and split accordingly in 2 groups: [1] those that are shared between target lineage, and [2] tumor-specific, not differentially expressed in cerebellum cell types. Gene ontology analysis was performed on selected tumor-specific groups using hypergeometric test on annotations from MSigDB reference database (*gsea-msigdb.org*) blocks C2 (curated gene sets) and C5 (ontology gene sets) with minimum p-value limit 0.01. Additional selection was performed from top50 significant terms from each gene ontology test.

For above detected genes we also inspected global bulk tissue profiles^3^ (n=297) to mark only brain-specific (obtained via differential expression analysis among tissues via limma package) for shared [1] and not/low expressed in any tissue (based on mean expression < 5 RPKM limit) for unique [2] as well as combined surfaceome markers and druggable candidates using Human Protein Atlas resource (*proteinatlas.org*).

### Single cell tumor transcriptome data processing

Single cell datasets were collated from various publications for medulloblastoma, ependymoma and PA as described in Suppl. Table2, representing 10x (v2 and v3) and SmartSeq2 platforms. SmartSeq2 sample data processing: Gene count matrices filtered for cells were normalized using *scran*^*57*^, except for PA samples where TPM normalized gene matrix was used. Mitochondrial, ribosomal (prefixes: RPS, RPL, MRPS or MRP) and cell cycle associated genes^56^ were removed before calling HVGs, and top 2000 HVGs were used to perform PCA. First 15 PCAs were used to cluster cells in individual samples.

10x sample data processing: Filtered gene count matrices were processed with additional cell filtering performed for mitochondrial (prefix: MT-) gene proportion (cutoff: 3 MAD above mean) and number of genes detected (> 200 and <6000) for all samples, except for Gillen et al. data. Count matrix were normalized using *scran* and processed similar to SmartSeq2 samples.

### Cell type assignment to tumor cells

SingleR method^31^ was modified with a Support Vector Machine (SVM) based fine tuning step to assign each tumor cell an identity to the cerebellum cell state annotation reference. In the first round of classification, *scran* normalized count matrix were used for both reference (cerebellum data) and target (tumor sample). Genes were subsetted to an intersection of top 2000 HVGs across tumor types and all the HVGs in cerebellum (*getTOPhvg* function, *scran*). Since tumor cells are enriched for a particular lineage (e.g.: SHH-MB enriched for GC lineage), to improve classification of non-tumor cells in tumor samples, we merged top 2000 HVGs across medulloblastoma, ependymoma and PA samples. A Spearman correlation was calculated for each test cell and each cell state in cerebellum reference using *SingleR* (method= wilcox, de=15) and top candidate matches were selected after filtering for one standard deviation from the highest correlation score.

For pruning, a calibrated SVM model generated with LinearSVC (max_iter=100000, class_weight = ‘balanced’, cv=10, *scikit-learn*, v1.0.1) was used for each cell. Tumor and cerebellum data were further cosine normalized, for genes used to calculate correlation in *SingleR*, to account for sample specific size factors. For SVM prediction of each cell, the cerebellum reference was further subsetted to cells associated with top candidate cell state annotations. The SVM model assigned probability value to each candidate cell state with the highest probability assigned as “best match” identity. All best match identities with >=50% probability were assigned to “pruned identity”. In case the probability of assignment was <50%, the pruned identity was assigned as NA (not assigned).

### Combination of single cell tumor data with cerebellum reference

To study the trajectory of tumor *w.r.t* to cerebellum cell type trajectory, we integrated each tumor sample to the best match cell type lineage of that sample’s tumor type (medulloblastoma: GC/UBC, ependymoma: Astrocytes and PA: Oligodendrocytes). We used batch reduction method to integrate tumor sample to the matching normal lineage, with the assumption that tumorigenic effect on cell transcriptome is independent (and hence orthogonal) to normal developmental program and can be “corrected”. Batch correction via MNN^58^ adjusts a cell’s expression component along the average batch (in our case, tumor) vector. Hence to avoid miscalculating tumor vector, in each tumor sample, we removed probable (best match identity as assigned by SVM) non-tumor cells (non-neuronal cell for medulloblastoma, non-astroglial cells for ependymoma and non-oligodendrocytic cells for PA). Verification of non-tumor cells was performed via copy number profiling of EPN tumor samples performed with InferCNV^56^ v1.3.2. We then recalculated HVGs for the filtered cells and used top 500 genes after intersection with top 5000 HVGs in the lineage. The reference and target data were cosine normalized in the subsetted gene space, merged and factorized using NMF (max_iter=1000000, init = ‘nndsvda’, solver = ‘mu’, random_state = 5; *scikit-learn* v1.0.1) to obtain 100 NMF factors for the merged data. The reduced data was then batch corrected for tumor status variable using *reducedMNN* (R package *batchelor*) to obtain corrected factors which were further used for UMAP visualization and diffusion map analysis (*DiffusionMap* function, R package *destiny*^*59*^).

In diffusion map analysis, a pseudotime (*dpt* function) value was calculated for each cell by averaging *dpt* values obtained from 30 randomly selected set of roots cells in reference data (Stage =CS18, cell subtype: medulloblastoma - early RL; ependymoma/PA - early VZ). For additional inspection of PA to compare a single cell tumor sample with the lineage of origin, the tumor single cell data, without any filtering for non-tumor cells, was combined with cells assigned to lineage of origin (e.g. oligodendrocytes). The gene space was nested to a union of top 1000 HVG from the tumor sample and lineage of origin. Resulting gene expression matrix was scaled using cosineNorm, combined and factorized using NMF as described above to obtain 100 NMF factors. The factorized data was used to obtain UMAP visualization of the combined tumor sample and normal cell type lineage.

### Available resources

The results of all described main comparisons of normal cell types to tumors (both bulk and single cell data) can be checked from online ShinyApp R application at *brain-match.org*. Source code of the application is available upon request.

## Notes

http://www.brain-match.org/

### Competing Interest Statement

The authors have declared no competing interest.

